# Single-cell RNA-seq of mouse dopaminergic neurons informs candidate gene selection for sporadic Parkinson’s disease

**DOI:** 10.1101/148049

**Authors:** Paul W. Hook, Sarah A. McClymont, Gabrielle H. Cannon, William D. Law, A. Jennifer Morton, Loyal A. Goff, Andrew S. McCallion

## Abstract

Genetic variation modulating risk of sporadic Parkinson’s disease (PD) has been primarily explored through genome wide association studies (GWAS). However, like many other common genetic diseases, the impacted genes remain largely unknown. Here, we used single-cell RNA-seq to characterize dopaminergic (DA) neuron populations in the mouse brain at embryonic and early postnatal timepoints. These data facilitated unbiased identification of DA neuron subpopulations through their unique transcriptional profiles, including a novel postnatal neuroblast population and *substantia nigra* (SN) DA neurons. We use these population-specific data to develop a scoring system to prioritize candidate genes in all 49 GWAS intervals implicated in PD risk, including known PD genes and many with extensive supporting literature. As proof of principle, we confirm that the nigrostriatal pathway is compromised in *Cplx1* null mice. Ultimately, this systematic approach establishes biologically pertinent candidates and testable hypotheses for sporadic PD, informing a new era of PD genetic research.

---

The most commonly used genetic tool today for studying complex disease is the genome wide association study (GWAS). As a strategy, GWAS was initially hailed for the insight it might provide into the genetic architecture of common human disease risk. Indeed, the collective data from GWAS since 2005 has revealed a trove of variants and genomic intervals associated with an array of phenotypes^1^. The majority of variants identified in GWAS are located in non-coding DNA^2^ and are enriched for characteristics denoting regulatory DNA^2,3^. This regulatory variation is expected to impact expression of a nearby gene, leading to disease susceptibility.

Traditionally, the gene closest to the lead SNP has been prioritized as the affected gene. However, recent studies show that disease-associated variants can act on more distally located genes, invalidating genes that were previously extensively studied^4,5^. The inability to systematically connect common variation with the genes impacted limits our capacity to elucidate potential therapeutic targets and can waste valuable research efforts.

Although GWAS is inherently agnostic to the context in which disease-risk variation acts, the biological impact of common functional variation has been shown to be cell context dependent^2,6^. Extending these observations, Pritchard and colleagues recently demonstrated that although genes need only to be expressed in disease-relevant cell types to contribute to risk, those expressed preferentially or exclusively therein contribute more per SNP^7^. Thus, accounting for the cellular and gene regulatory network (GRN) contexts within which variation act may better inform the identification of impacted genes. These principles have not yet been applied systematically to many of the traits for which GWAS data exists. We have chosen Parkinson’s disease (PD) as a model complex disorder for which a significant body of GWAS data remains to be explored biologically in a context dependent manner.

PD is the most common progressive neurodegenerative movement disorder. Incidence of PD increases with age, affecting an estimated 1% worldwide beyond 70 years of age^8–10^. The genetic underpinnings of non-familial or sporadic PD have been studied through the use of GWAS with recent meta-analyses highlighting 49 loci associated with sporadic PD susceptibility^11,12^. While a small fraction of PD GWAS loci contain genes known to be mutated in familial PD (*SNCA* and *LRRK2*)^13,14^, most indicted intervals do not contain a known causal gene or genes. Although PD ultimately affects multiple neuronal centers, preferential degeneration of DA neurons in the SN leads to functional collapse of the nigrostriatal pathway and loss of fine motor control. The preferential degeneration of SN DA neurons in relation to other mesencephalic DA neurons has driven research interest in the genetic basis of selective SN vulnerability in PD. Consequently, one can reasonably assert that a significant fraction of PD-associated variation likely mediates its influence specifically within the SN.

In an effort to illuminate a biological context in which PD GWAS results could be better interpreted, we undertook single-cell RNA-seq (scRNA-seq) analyses of multiple DA neuronal populations in the brain, including ventral midbrain DA neurons. This analysis defined the heterogeneity of DA populations over developmental time in the brain, revealing gene expression profiles specific to discrete DA neuron subtypes. These data further facilitated the definition of GRNs active in DA neuron populations including the SN. With these data, we establish a framework to systematically prioritize candidate genes in all 49 PD GWAS loci and begin exploring their pathological significance.

## RESULTS

### scRNA-seq characterization defines DA neuronal subpopulation heterogeneity

In order to characterize DA neuron molecular phenotypes, we undertook scRNA-seq on cells isolated from distinct anatomical locations of the mouse brain over developmental time. We used fluorescence activated cell sorting (FACS) to retrieve single DA neurons from the Tg(Th-EGFP)DJ76Gsat BAC transgenic mouse line, which expresses eGFP under the control of the tyrosine hydroxylase (*Th*) locus^15^. We microdissected both MB and FB from E15.5 mice, extending our analyses to MB, FB, and OB in P7 mice (Figure 1a). E15.5 and P7 time points were chosen based on their representation of stable MB DA populations, either after neuron birth (E15.5) or between periods of programmed cell death (P7) (Figure 1a)^16^.

**Figure 1.**
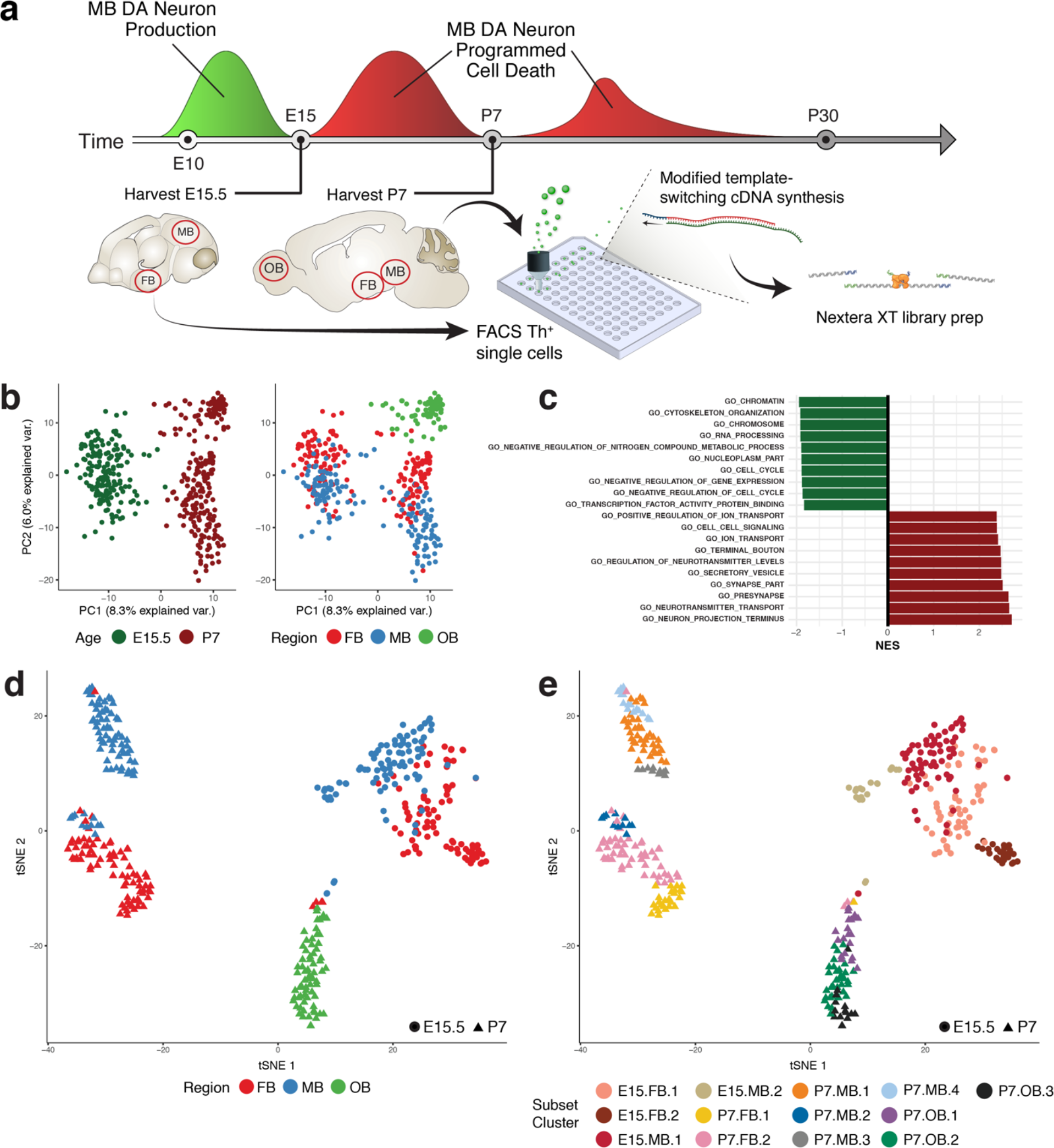
scRNA-seq analysis of isolated cells allows their separation by developmental time. Figure 1. scRNA-seq analysis of isolated cells allows their separation by developmental time. a) Diagram of scRNA-seq experimental procedures for isolating and sequencing EGFP+ cells. Timeline adapted from Barallobre, et al., 2014a. b) Principal component analysis (PCA) on all cells collected using genes with highly variant transcriptional profiles. The greatest source of variation (PC1) is explained by the time point at which the cells were collected, not the region from which the cells were collected. c) The top ten Gene Ontology (GO) gene sets enriched in genes with positive (red) and negative (green) PC1 loadings. Genes with negative PC1 loadings and negative normalized enrichment scores (NES) were enriched for terms indicative of mitotically active cells. Genes with positive PC1 loadings and NES scores were enriched for terms expected of more mature neurons. d) A t-distributed Stochastic Neighbor Embedding (t-SNE) plot of all collected cells colored by regional identity. E15.5 cells cluster together while P7 cells cluster primarily by regional identity. e) A t-SNE plot of all collected cells colored by subset cluster identity. Through iterative analysis, timepoint-regions collected can be separated into multiple subpopulations (13 in total). Midbrain, Mb; Forebrain, FB; Olfactory bulb; OB; Fluorescence activated cell sorting; FACS.

Quality control and outlier analysis identify 396 high quality cell transcriptomes to be used in our analyses. We initially sequenced RNA from 473 single cells to an average depth of ~8 × 10^5^ 50 bp paired-end fragments per cell. Using Monocle 2, we converted normalized expression estimates into estimates of RNA copies per cell^17^. Cells were filtered based on the distributions of total mass, total number of mRNAs, and total number of expressed genes per cell (Figure 1 - figure supplement 1a-1c; detailed in Methods). After QC, 410 out of 473 cells were retained. Using principal component analysis (PCA) as part of the iterative analysis described below, we identified and removed 14 outliers determined to be astrocytes, microglia, or oligodendrocytes (Figure 1 - figure supplement 1e; Supplementary File 1), leaving 396 cells (~79 cells/timepoint-region; Figure 1 - figure supplement 1d).

**Figure 1 - supplement 1.**
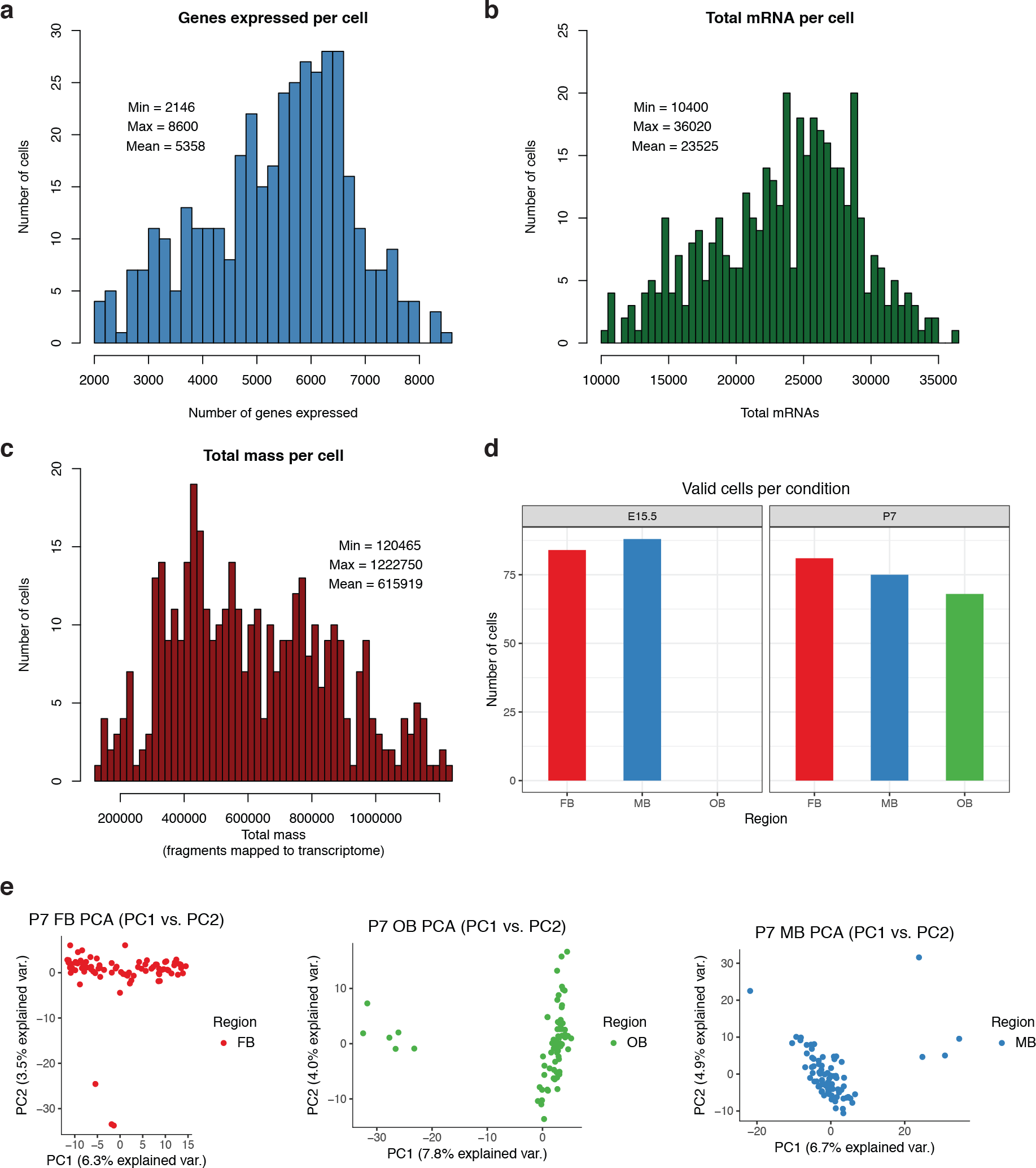
Quality control used for filtering single-cell RNA-seq data. Figure 1 - supplement 1. Quality control used for filtering single-cell RNA-seq data. a) Histogram showing the final distribution of the number of genes expressed per cell (n cells = 396). b) Histogram showing the final distribution of the total mRNA per cell (n cells = 396). c) Histogram showing the final distribution of the total mass (fragments mapped to the transcriptome) per cell (n cells = 396). d) Barplot showing the number of cells in each timepoint-region. There were a mean of 79 cells/timepoint region. e) Principal component analysis (PCA) plots from the iterative analyses performed on P7 FB, P7 OB, and P7 MB cell populations. Initial analyses in these timepoint-regions revealed outliers that were subsequently removed.

**Figure 1 - supplement 2.**
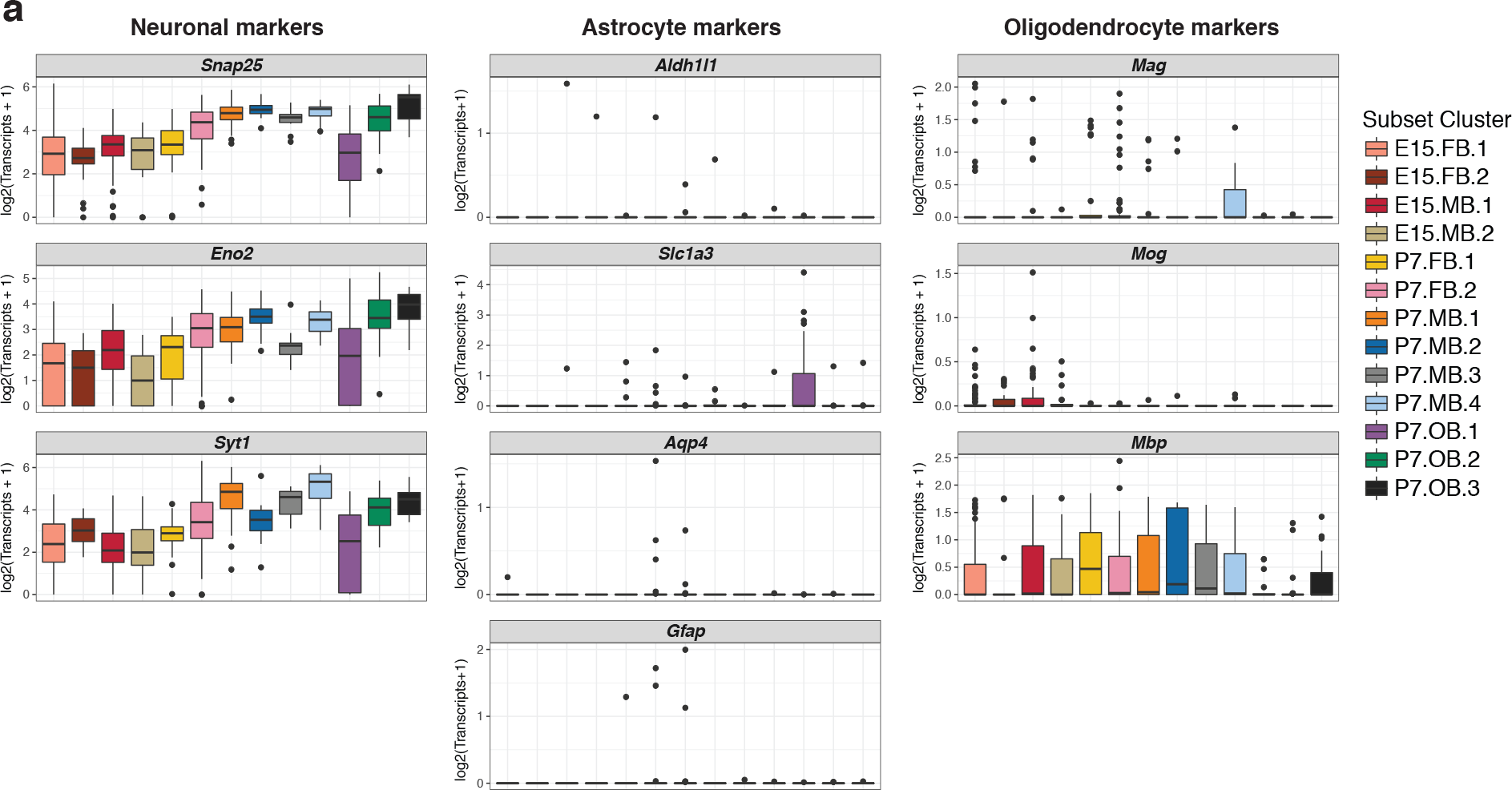
Expression of broad marker genes confirms successful isolation of neurons. Figure 1 - supplement 2. Expression of broad marker genes confirms successful isolation of neurons. a) Boxplots showing the expression of pan-neuronal, pan-astrocyte, and pan-oligodendrocyte marker in all 13 subpopulations. All subpopulations show robust expression of pan-neuronal markers. +/− 1.5× interquartile range is represented by the whiskers on the boxplots. Data points beyond 1.5× interquartile range are considered as outliers and plotted as black points.

To confirm that our methods can discriminate between different populations of neurons, we first explored differences between timepoints. Following a workflow similar to the recently described “dpFeature” procedure^18^, we identified genes with highly variable transcriptional profiles and performed PCA. As anticipated, we observed that the greatest source of variation was between developmental ages (Figure 1b). Genes associated with negative PC1 loadings (E15.5 cells) were enriched for gene sets consistent with mitotically active neuronal, undifferentiated precursors (Figure 1c). In contrast, genes associated with positive PC1 loadings (P7 cells) were enriched for ontology terms associated with mature, post-mitotic neurons (Figure 1c). This initial analysis establishes our capacity to discriminate among biological classes present in our data using PCA as a foundation.

Further, we attempted to identify clusters of single cells between and within timepoints and anatomical regions. In order to do this, we selected the PCs that described the most variance in the data and used t-Stochastic Neighbor Embedding (t-SNE)^19^ to further cluster cells in an unsupervised manner (see Methods). Analysis of all cells revealed that the E15.5 cells from both MB and FB cluster together (Figure 1d), supporting the notion that they are less differentiated. By contrast, cells isolated at P7 mostly cluster by anatomical region, suggesting progressive functional divergence with time (Figure 1d). We next applied this same scRNA-seq analysis workflow (See Methods) in a recursive manner individually in all regions at both timepoints to further explore heterogeneity. This revealed a total of 13 clusters (E15.5 FB.1-2, MB.1-2; P7 OB.1-3, FB.1-2, MB.1-4; Figure 1e), demonstrating the diversity of DA neuron subtypes and providing a framework upon which to evaluate the biological context of genetic association signals across closely-related cell types. Using known markers, we confirmed that all clusters expressed high levels of pan-neuronal markers (*Snap25*, *Eno2*, and *Sytl*) (Figure 1 - figure supplement 2a). In contrast, we observed scant evidence of astrocyte (*Aldh1l1*, *Slc1a3*, *Aqp4*, and *Gfap*; Figure 1 - figure supplement 2a) or oligodendrocyte markers *(Mag, Mog,* and *Mbp*; Figure 1 - figure supplement 2a), thus confirming we successfully isolated our intended substrate, *Th*+ neurons.

**Figure 2.**
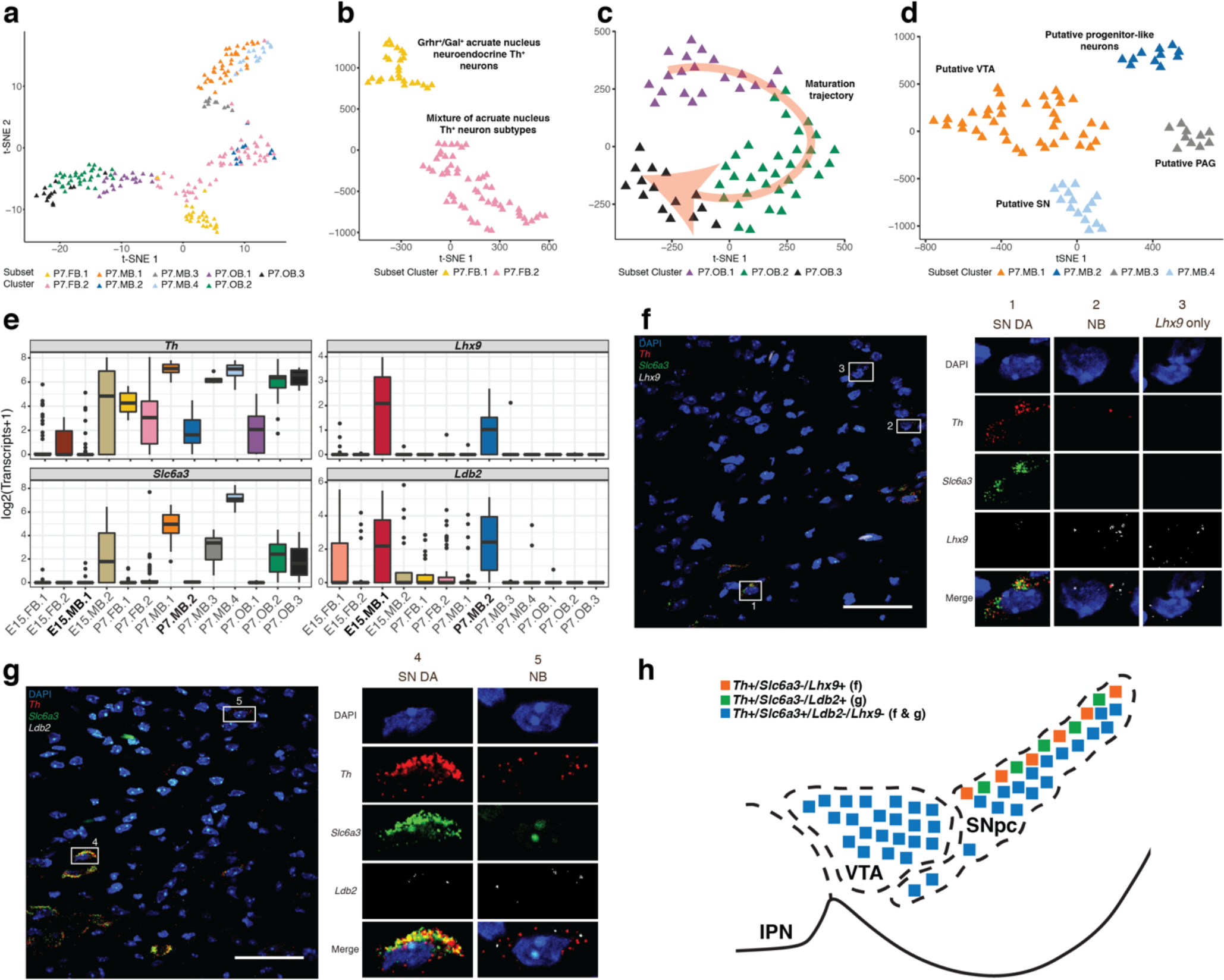
Subclusters of P7 *Th*+ neurons are identified based on marker gene analyses. Figure 2. Subclusters of P7 *Th*+ neurons are identified based on marker gene analyses. a) A t-SNE plot of all P7 neurons collected using colored by subset cluster identity. The neurons mostly cluster by regional identity. b) t-SNE plot of P7 FB neurons. P7 FB neurons cluster into two distinct populations. c) t-SNE plot of P7 OB neurons. P7 OB neurons cluster into three populations. These populations represent a trajectory of Th+ OB maturation (Table S3) as indicated by the red arrow. d) A t-SNE plot of P7 MB neurons. P7 MB neurons cluster into four clusters: the *substantia nigra* (SN), the ventral tegmental area (VTA), the periaqueductal grey area (PAG), and a novel progenitor-like population. e) Boxplots displaying the expression of four genes (*Th*, *Slc6a3*, *Lhx9*, and *Ldb2*) across all subclusters identified. The novel P7 MB progenitor-like cluster (P7.MB.2) has a similar expression profile to E15.5 MB neuroblast population (E15.MB. 1) (Table S2). +/− 1.5× interquartile range is represented by the whiskers on the boxplots. Data points beyond 1.5× interquartile range are considered as outliers and plotted as black points. f) Representative image of multiplex single molecule fluorescent *in situ* hybridization (smFISH) for *Th*, *Slc6a3*, and *Lhx9*, in the mouse ventral midbrain. Zoomed-in panels represent cell populations observed. Scale bar, 50 uM. g) Representative image of multiplex smFISH for *Th*, *Slc6a3*, and *Ldb2*, in the mouse ventral midbrain. Zoomed-in panels represent cell populations observed. h) Diagram of ventral midbrain summarizing the results of smFISH. Th+/Slc6a3-/Lhx9+ and Th+/Slc6a3-/Ldb2+ cells are both found in the dorsal SN. Scale bar, 50 uM. NB, neuroblast; SN, substantia nigra; VTA, ventral tegmental area; IPN, interpeduncular nucleus.

**Figure 2 - supplement 1.**
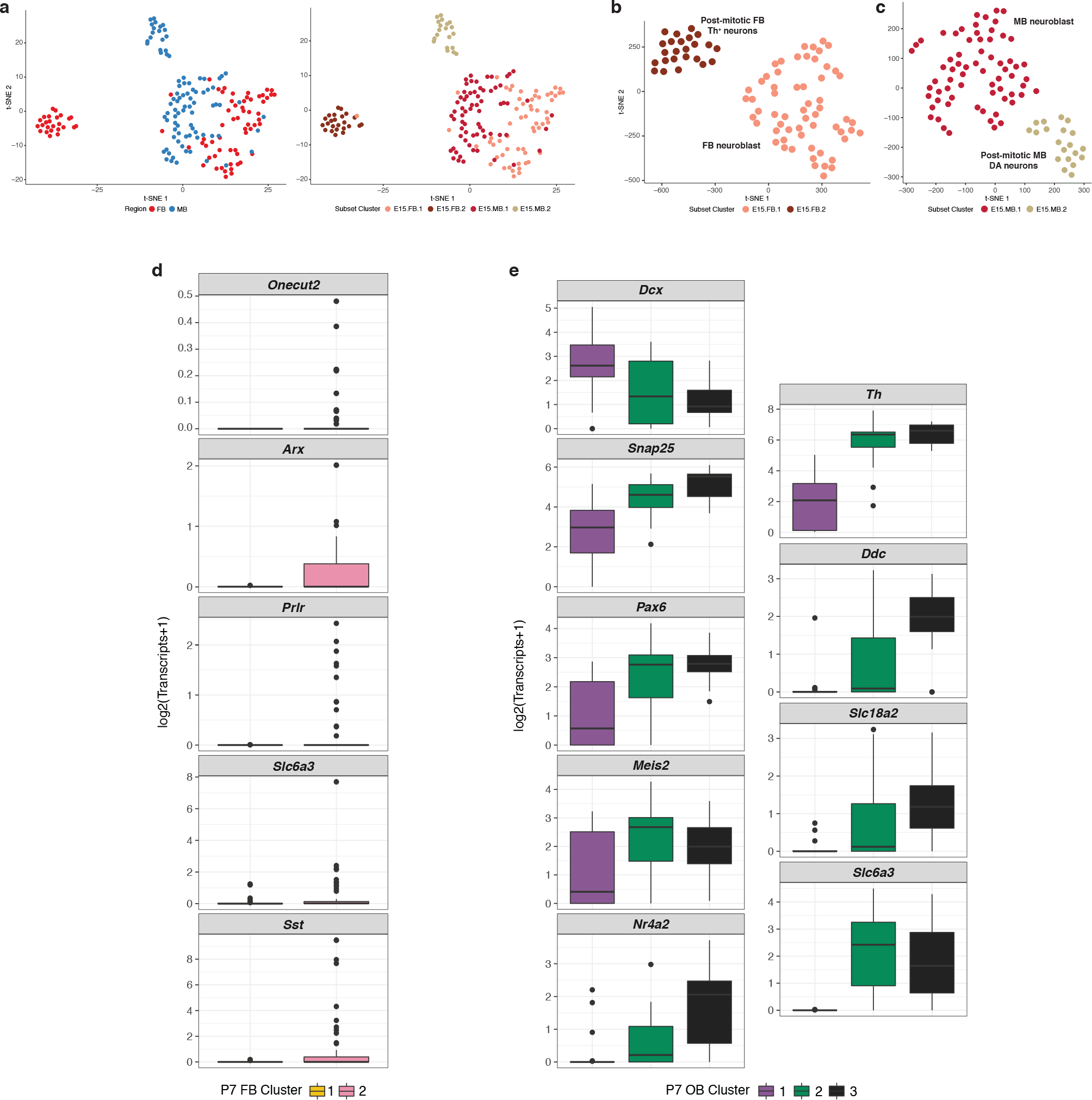
Clusters of Th+ neurons are discovered through iterative, marker gene analysis. Figure 2 - supplement 2. Clusters of Th+ neurons are discovered through iterative, marker gene analysis. a) t-SNE plots of all E15.5 cells colored by regional identity and subset cluster assignment. b) t-SNE plot of FB E15.5 cells colored by subset cluster assignment. E15.5 FB cells cluster in two distinct populations. c) t-SNE plot of MB E15.5 cells colored by subset cluster assignment. E15.5 MB cells cluster in two distinct populations. d) Boxplots showing the expression of markers used to identify the P7.FB.2 cluster (Table S3). +/− 1.5× interquartile range is represented by the whiskers on the boxplots. Data points beyond 1.5× interquartile range are considered as outliers and plotted as black points. e) Boxplots showing the expression of markers used to identify P7 olfactory bulb clusters (Table S3). +/− 1.5× interquartile range is represented by the whiskers on the boxplots. Data points beyond 1.5× interquartile range are considered as outliers and plotted as black points.

### scRNA-seq revealed biologically and temporally discriminating transcriptional signatures

With subpopulations of DA neurons defined in our data, we set out to assign a biological identity to each cluster. Among the four clusters identified at E15.5, two were represented in t-SNE space as a single large group that included cells from both MB and FB (E15.MB.1, E15.FB.1), leaving two smaller clusters that were comprised solely of MB or FB cells (Figure 2 - figure supplement 1a). The latter MB cluster (E15.MB.2; Figure 2 - figure supplement 1a-1c) specifically expressed *Foxa1*, *Lmx1a*, *Pitx3*, and *Nr4a2* and thus likely represents a post-mitotic DA neuron population^20^ (Supplementary File 2; Supplementary File 3). Similarly, the discrete E15.FB.2 cluster expressed markers of post-mitotic FB/hypothalamic neurons (Figure 2 - figure supplement 1a-1b), including *Six3*, *Six3os1*, *Sst*,and *Npy* (Supplementary File 2; Supplementary File 3). These embryonic data did not discriminate between cells populating known domains of DA neurons, such as the SN.

By contrast, P7 cells mostly cluster by anatomical region and each region has defined subsets (Figure 1d, 1e, 2a). Analysis of P7 FB revealed two distinct cell clusters (Figure 2b). Expression of the neuropeptides *Gal* and *Ghrh* and the *Gsx1* transcription factor place P7.FB.1 cells in the arcuate nucleus (Supplementary File 2; Supplementary File 3)^21–24^. The identity of P7.FB.2, however, was less clear, although subsets of cells therein did express other arcuate nucleus markers for *Th*^+^/*Ghrh*^−^ neuronal populations e.g. *Onecut2*, *Arx*, *Prlr*, *Slc6a3*, and *Sst* (Figure 2 - figure supplement 1d; Supplementary File 3)^24^. All three identified OB clusters (Figure 2c) express marker genes of OB DA neuronal development or survival (Supplementary File 2, Supplementary File 3; Figure 2 - figure supplement 1e)^25^. It has previously been reported that *Dcx* expression diminishes with neuronal maturation^26^ and *Snap25* marks mature neurons^27^. We observe that these OB clusters seem to reflect this continuum of maturation wherein expression of *Dcx* diminishes and *Snap25* increases with progression from P7.OB1 to OB3 (Figure 2 - figure supplement 1e). This pattern is mirrored by a concomitant increase in OB DA neuron fate specification genes (Figure 2 - figure supplement 1e)^25,28^. In addition, we identified four P7 MB DA subset clusters (Figure 2d). Marker gene analysis confirmed that three of the clusters correspond to DA neurons from the VTA (*Otx2* and *Neurod6*; P7.MB.1)^29,30^, the PAG (*Vip* and *Pnoc*; P7.MB.3)^31,32^, and the SN (*Sox6*, *Aldh1a7*, *Ndnf*, *Serpine2*, *Rbp4*, and *Fgf20*; P7.MB.4)^29,33–35^ (Supplementary File 2; Supplementary File 3). These data are consistent with recent scRNA-seq studies of similar populations^34,36^. Through this marker gene analysis, we successfully assigned a biological identity to 12/13 clusters.

The only cluster without a readily assigned identity was P7.MB.2. This population of P7 MB DA neurons, P7.MB.2 (Figure 2d), is likely a progenitor-like population. Like the overlapping E15.MB.1 and E15.FB.1 clusters (Figure 2 - figure supplement 1a), this cluster preferentially expresses markers of neuronal precursors/differentiation/maturation (Supplementary File 2, Supplementary File 3). In addition to sharing markers with the progenitor-like E15.MB.1 cluster, P7.MB.2 exhibits gene expression consistent with embryonic mouse neuroblast populations^34^, cell division, and neuron development^37–41^ (Supplementary File 2, Supplementary File 3). Consistent with the hypothesis, this population displayed lower levels of both *Th* and *Slc6a3*, markers of mature DA neurons, than the terminally differentiated and phenotypically discrete P7 MB DA neuron populations of the VTA, SN and PAG (Figure 2e).

With this hypothesis in mind, we sought to ascertain the spatial distribution of P7.MB.2 DA neurons through multiplex, single molecule fluorescence *in situ* hybridization (smFISH) for *Th* (pan-P7 MB DA neurons), *Slc6a3* (P7.MB.1, P7.MB.3, P7.MB.4), and one of the neuroblast marker genes identified through our analysis, either *Lhx9* or *Ldb2* (P7.MB.2) (Figure 2e). In each experiment, we scanned the ventral midbrain for cells that were Th+/Slc6a3- and positive for the third gene. *Th*+/*Slc6a3*-/*Lhx9*+ cells were found scattered in the dorsal SN *pars compacta* (SNpc) along with cells expressing *Lhx9* alone (Figure 2f, 2h). Expression of *Ldb2* was found to have a similar pattern to *Lhx9,* with *Th*+/*Slc6a3*-/*Ldb2*+ cells found in the dorsal SNpc (Figure 2f, 2h). Expression of *Lhx9* and *Ldb2* was low or non-existent in *Th+/Slc6a3+* cells in the SNpc (Figure 2e, 2f). Importantly, cells expressing these markers express *Th* at lower levels than *Th*+/*Slc6a3*+ neurons (Figure 2f, 2g), consistent with our scRNA-seq data (Figure 2e). Thus, with the resolution of the spatial distribution of this novel neuroblast-like P7 MB DA population, we assign biological identity to each defined brain DA subpopulation.

### Novel SN-specific transcriptional profiles and GRNs highlight its association with PD

Overall our analyses above allowed us to successfully separate and identify 13 brain DA neuronal populations present at E15.5 and P7, including SN DA neurons. Motivated by the clinical relevance of SN DA neurons to PD, we set out to understand what makes them transcriptionally distinct from the other MB DA neuron populations.

In order to look broadly at neuronal subtypes, we evaluated expression of canonical markers of other neuronal subtypes in our *Th*+ neuron subpopulations. Interestingly, we observed inconsistent detection of *Th* and eGFP in some E15.5 clusters (Figure 3 - figure supplement 1a). This likely reflects lower *Th* transcript abundance at this developmental state, but sufficient expression of the eGFP reporter to permit FACS collection (Figure 3 - figure supplement 1b). The expression of other DA markers, *Ddc* and *Slc18a2*, mirror *Th* expression, while *Slc6a3* expression is more spatially and temporally restricted (Figure 3 - figure supplement 1a). The SN cluster displays robust expression of all canonical DA markers (Figure 3 - figure supplement 1a). Multiple studies have demonstrated that *Th*+ neurons may also express markers characteristic of other major neuronal subtypes^42–44^. We found that only the SN and PAG showed no expression of either GABAergic (*Gad1/Gad2/Slc32a1*) or glutamatergic (*Slc17a6*) markers (Figure 3 - figure supplement 1a). This neurotransmitter specificity is a potential avenue for exploring the preferential vulnerability of the SN in PD.

**Figure 3.**
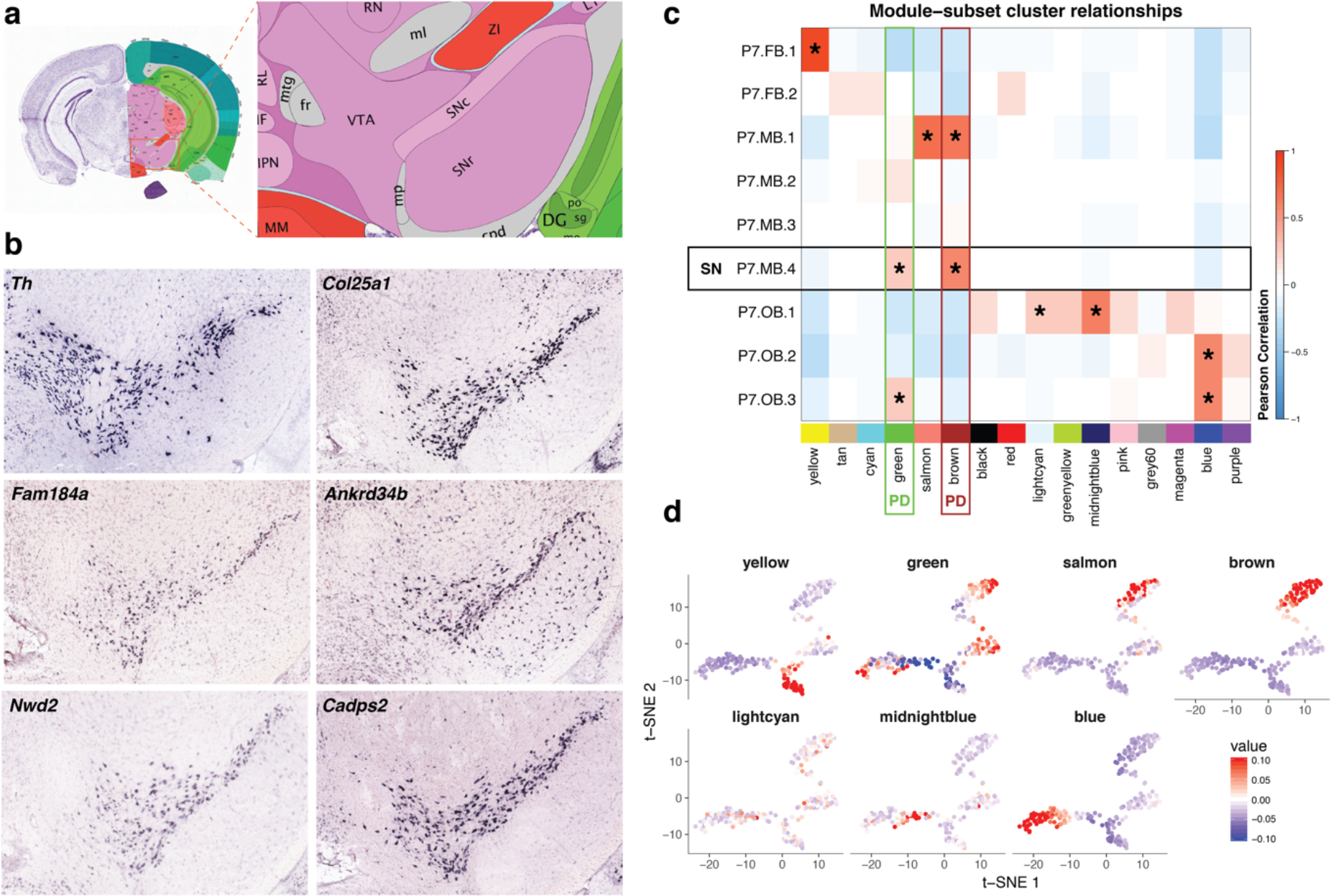
Novel markers and gene modules reveal context specific SN DA biology. Figure 3. Novel markers and gene modules reveal context specific SN DA biology. a) Reference atlas diagram from the Allen Brain Atlas (ABA; http://www.brain-map.org/) of the P56 mouse ventral midbrain. b) Confirmation of novel SN DA neuron marker genes through the use of ABA *in situ* hybridization data (http://www.brain-map.org/). Coronal, P56 mouse *in situ* data was explored in order to confirm the expression of 25 novel SN markers. *Th* expression in P56 mice was used as an anatomical reference during analysis. c) Correlation heatmap of the Pearson correlation between module eigengenes and P7 Th+ subset cluster identity. Modules are represented by their assigned colors at the bottom of the matrix. Modules that had a positive correlation with a subset cluster and had a correlation P-value less than the Bonferroni corrected significance level (P-value < 3.5e-04) contain an asterisk. SN cluster (P7.MB.4) identity is denoted by a black rectangle. Modules (“green” and “brown”) that were enriched for the “Parkinson’s Disease” KEGG gene set are labeled with “PD.” d) The eigengene value for each P7 neuron in the seven WGCNA modules shown to be significantly positively associated with a subset cluster overlaid on the t-SNE plot of all P7 neurons (Figure 2a). Plotting of eigengenes confirms strict spatial restriction of module association. Only the “lightcyan” module does not seem to show robust spatial restriction.

**Figure 3 - supplement 1.**
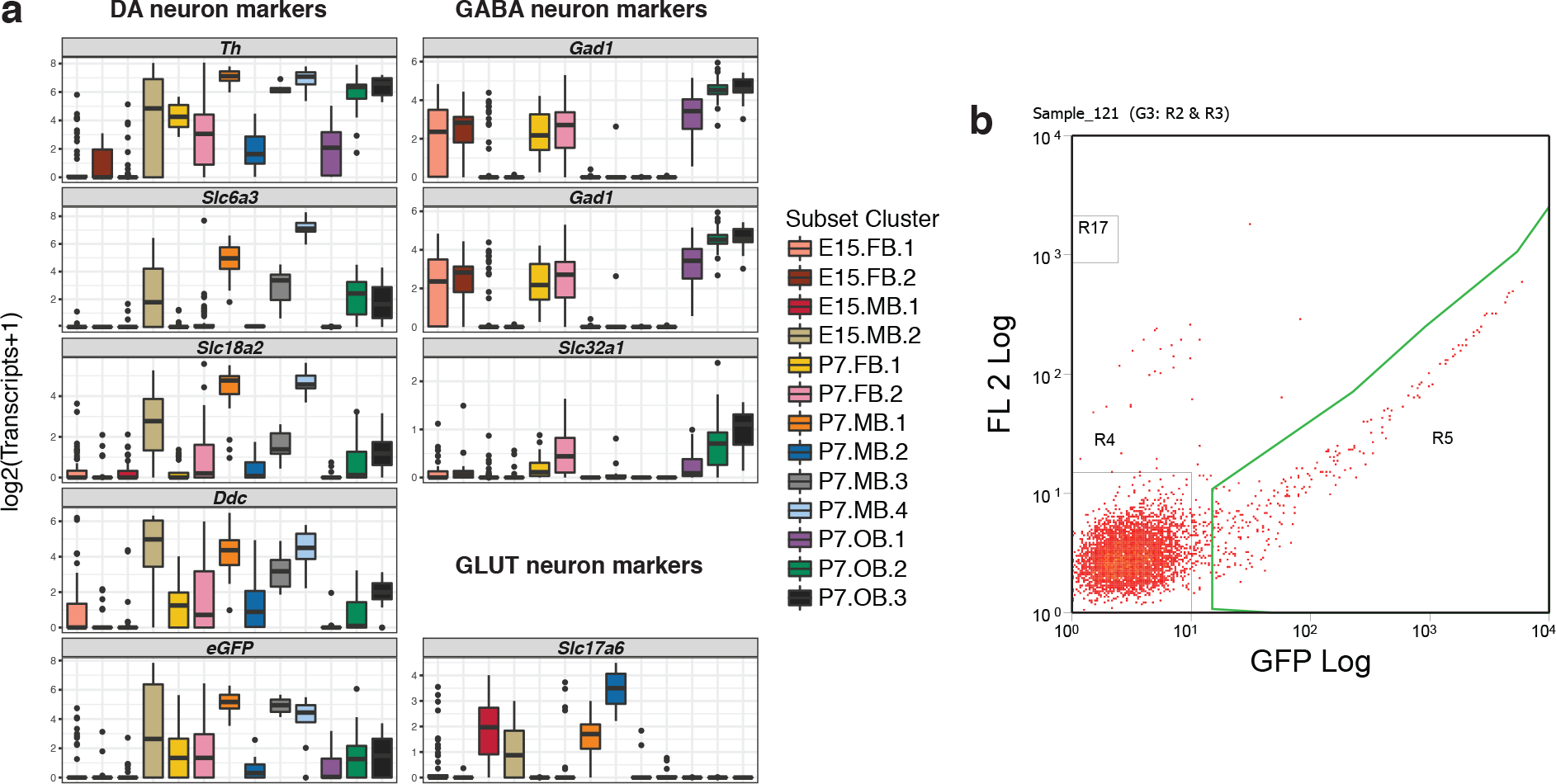
Exploration of neuronal subtype markers in isolated DA neuron populations. Figure 3 - supplement 1. Exploration of neuronal subtype markers in isolated DA neuron populations. a) Boxplots showing the expression of markers for dopaminergic (DA), GABAergic, or glutamatergic neurons. +/− 1.5× interquartile range is represented by the whiskers on the boxplots. Data points beyond 1.5× interquartile range are considered as outliers and plotted as black points. b) Example of a fluorescence activated cell sorting (FACS) plot used to isolate EGFP+ cells. EGFP fluorescence levels are represented on the x-axis and RFP fluorescence levels are represented on the y-axis. Cells were collected that fell within the area outlined in green.

**Figure 3 - supplement 2.**
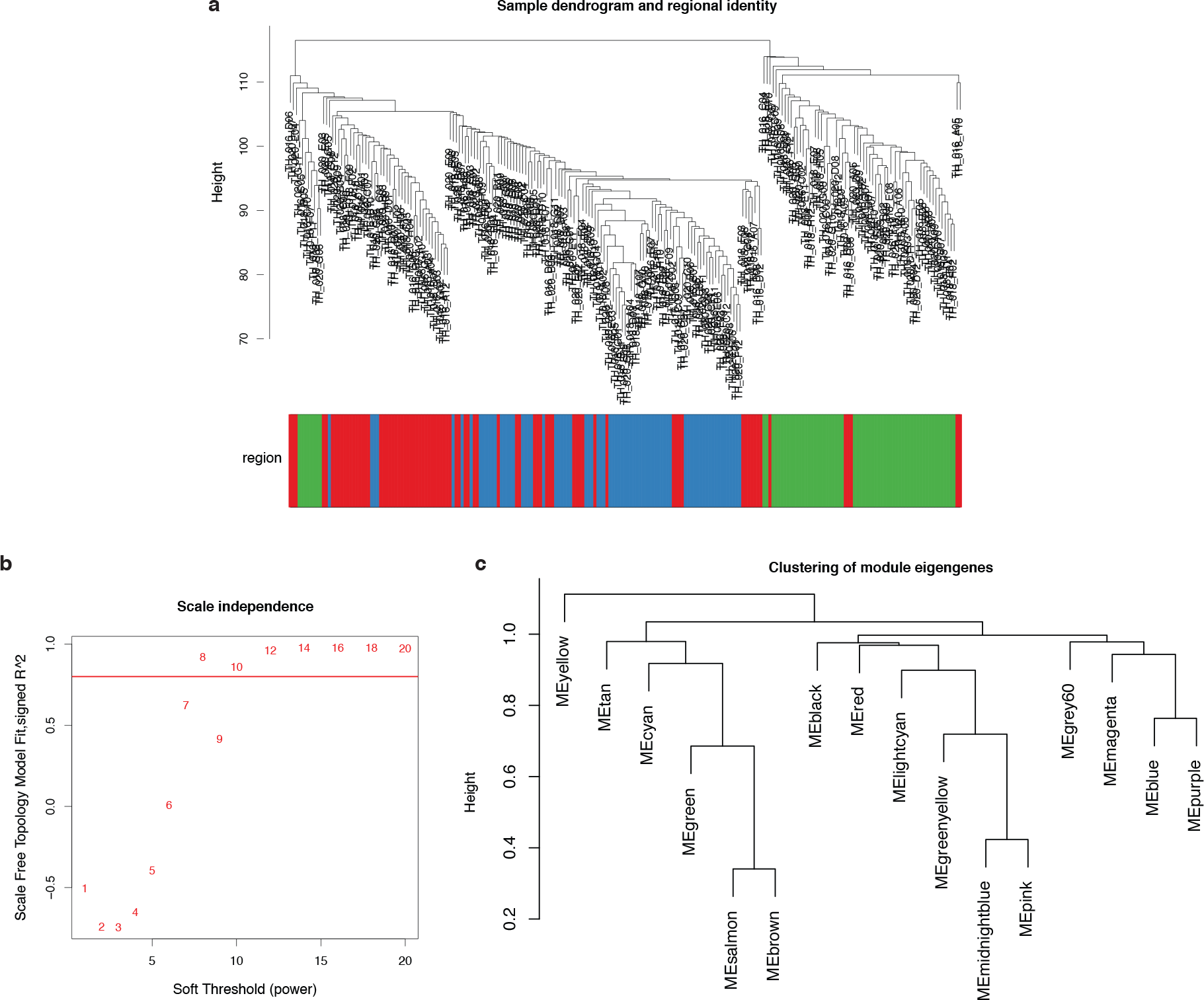
WGCNA analysis reveals 16 modules in P7 scRNA-seq data. Figure 3 - supplement 2. WGCNA analysis reveals 16 modules in P7 scRNA-seq data. a) A dendrogram of showing the relationship of P7 cells based on expressed genes. The cells are annotated by regional identity. b) Scale independence plot showing the scale free topology model fit for different levels of soft threshold power. This plot was used to determine the soft threshold that would be used for the rest of the analysis (soft threshold = 10). c) Hierarchical clustering shows the relationship between identified WGCNA modules.

Next, we postulated that genes whose expression defined the P7 SN DA neuron cluster might illuminate their preferential vulnerability in PD. We identified 110 SN-specific genes, by first finding all differentially expressed genes between P7 subset clusters and then using the Jensen-Shannon distance to identify cluster specific genes (See Methods). Prior reports confirm the expression of 49 of the 110 SN-specific genes (~45%) in postnatal SN (Supplementary File 4). We then sought evidence to confirm or exclude SN expression for the remaining, novel 61 genes (55%). Of these, 25/61 (~41%) were detected in adult SN neurons by *in situ* hybridization (ISH) of coronal sections in adult (P56) mice (Allen Brain Atlas, ABA; http://developingmouse.brain-map.org), including *Col25a1*, *Fam184a*, *Ankrd34b*, *Nwd2*, and *Cadps2* (Figure 3a, Supplementary File 5). Only 4/61 genes, for which ISH data existed in the ABA, lacked clear evidence of expression in the adult SN (Supplementary File 5). The ABA lacked coronal ISH data on 32/61 genes, thus we were unable to confirm their presence in the SN. Collectively, we identify 110 postnatal SN DA marker genes and confirm the expression of those genes in the adult mouse SN for 74 (67%) of them, including 25 novel markers of this clinically relevant cell population that we confirmed using the ABA image catalog.

We next asked whether we could identify significant relationships between cells defined as being P7 SN DA neurons and distinctive transcriptional signatures in our data. We identify 16 coexpressed gene modules by performing weighted gene co-expression network analysis (WGCNA)^45,46^ on all expressed genes of the P7 subset (Figure 3 - figure supplement 2; Supplementary File 6). By calculating pairwise correlations between modules and P7 subset clusters, we reveal that 7/16 modules are significantly and positively correlated (Bonferroni corrected p < 3.5e-04) with at least one subset cluster (Figure 3c). We graphically represent the eigenvalues for each module in each cell in P7 t-SNE space, confirming that a majority of these significant modules (6/7) displayed robust spatial, isotype enrichment (Figure 3d).

In order identify the biological relevance of these modules, each module was tested for enrichment for Kyoto Encyclopedia of Genes and Genomes (KEGG) pathways, Gene Ontology (GO) gene sets, and Reactome gene sets. Two modules, the “brown” and “green” modules, were significantly associated with the Parkinson’s Disease KEGG pathway gene set (Figure 3c; Supplementary File 7). Interestingly, the “brown” module was also significantly correlated with the P7 VTA population (P7.MB.1) and enriched for addiction gene sets (Supplementary File 7) highlighting the link between VTA DA neurons and addiction^47^. Strikingly, only the P7 SN cluster was significantly correlated with both PD-enriched modules (Figure 3c). This specific correlation suggests these gene modules may play a role in the preferential susceptibility of the SN in PD.

### Integrating SN DA neuron specific data enables prioritization of genes within PD-associated intervals

With these context-specific data in hand, we posited that SN DA neuron-specific genes and the broader gene co-expression networks that correlate with SN DA neurons might be used to prioritize genes within loci identified in PD GWAS. Such a strategy would be agnostic to prior biological evidence and independent of genic position relative to the lead SNP, the traditional method used to prioritize causative genes.

To investigate pertinent genes within PD GWAS loci, we identified all human genes within topologically associated domains (TADs) and a two megabase interval encompassing each PD-associated lead SNP. TADs were chosen because regulatory DNA impacted by GWAS variation is more likely to act on genes within their own TAD^48^. While topological data does not exist for SN DA neurons, we use TAD boundaries from hESCs as a proxy, as TADs are generally conserved across cell types^49^. To improve our analyses, we also selected +/- 1 megabase interval around each lead SNP thus including the upper bounds of reported enhancer-promoter interactions^50,51^. All PD GWAS SNPs interrogated were identified by the most recent metaanalyses (49 SNPs in total)^11,12^, implicating a total of 1751 unique genes. We then identified corresponding one-to-one mouse to human homologs (1009/1751; ~58%), primarily through the Mouse Genome Informatics (MGI) homology database.

To prioritize these genes in GWAS loci, we developed a gene-centric score that integrates our data as well as data in the public domain. We began by intersecting the PD loci genes with our scRNA-seq data as well as previously published SN DA expression data^34^, identifying 430 genes (430/1009; ~43%) with direct evidence of expression in SN DA neurons in at least one dataset. Each PD-associated interval contained ≥1 SN-expressed gene (Supplementary File 8). Emphasizing the need for a novel, systematic strategy, in 19/49 GWA intervals (~39%), the most proximal gene to the lead SNP was not detectably expressed in mouse SN DA neuron populations (Supplementary File 8; Supplementary File 9). Surprisingly, three loci contained only one SN DA-expressed gene: *Mmp16* (*MMP16* locus, Figure 4a), *Tsnax* (*SIPA1L2* locus), and *Satb1* (rs4073221 locus). The relevance of these candidate genes to neuronal function/dysfunction is well supported^52–56^. This establishes gene expression in a relevant tissue as a powerful tool in the identification of causal genes.

**Figure 4.**
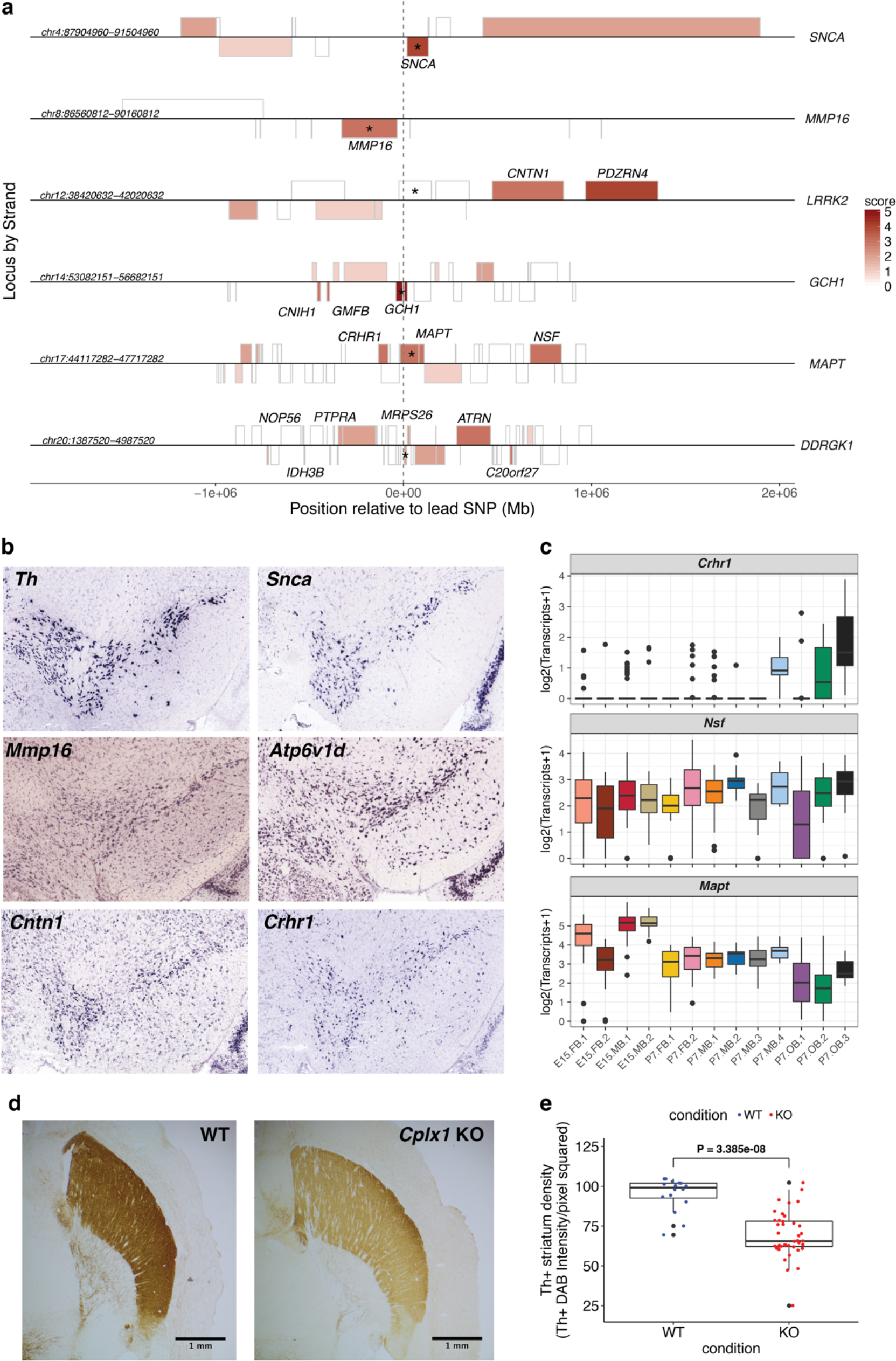
Context specific SN DA data allows for the prioritization of genes in PD GWAS loci. Figure 4. Context specific SN DA data allows for the prioritization of genes in PD GWAS loci. a) A locus plot displaying four megabase regions in the human genome (hg38) centered on PD GWAS SNPs in six loci. Genes are displayed as boxes on their appropriate strand. Genes are shaded by their prioritization score and gene names are displayed for genes with a score of 3 or higher in each locus. b) *In situ* hybridization from the ABA (http://www.brain-map.org/) of five prioritized genes along with *Th* for an anatomical reference. Coronal, P56 mouse *in situ* data was used. c) Boxplots displaying expression of prioritized genes from the *MAPT* locus (Figure 4a; Table 1). +/− 1.5× interquartile range is represented by the whiskers on the boxplots. Data points beyond 1.5× interquartile range are considered as outliers and plotted as black points. d) Representative light microscopy images of *Th*+ innervation density in the striatum of WT and *Cplx1* knockout (KO) mice. Scale bar, 1 mm. e) Boxplots comparing the level of Th+ striatum innervation between WT and *Cplx1* KO mice. DAB staining density was measured in 35 uM, horizontal sections in WT mice (mice = 3, sections = 16) and *Cplx1* KO mice (mice = 8, sections = 40). Each point in the boxplot represents a stained, 35 uM section. Statistical analyses were performed between conditions with section averages in order to preserve observed variability (WT n = 16, Cplx1 KO n = 40). A two sample t-test revealed that Th+ innervation density was significantly lower in *Cplx1* KO mice (t = 6.4395, df = 54, p = 3.386e-08). Data points outside of 1.5x interquartile range, represented by the whiskers on the boxplots, are considered as outliers and plotted as black points.

**Figure 4 - supplement 1.**
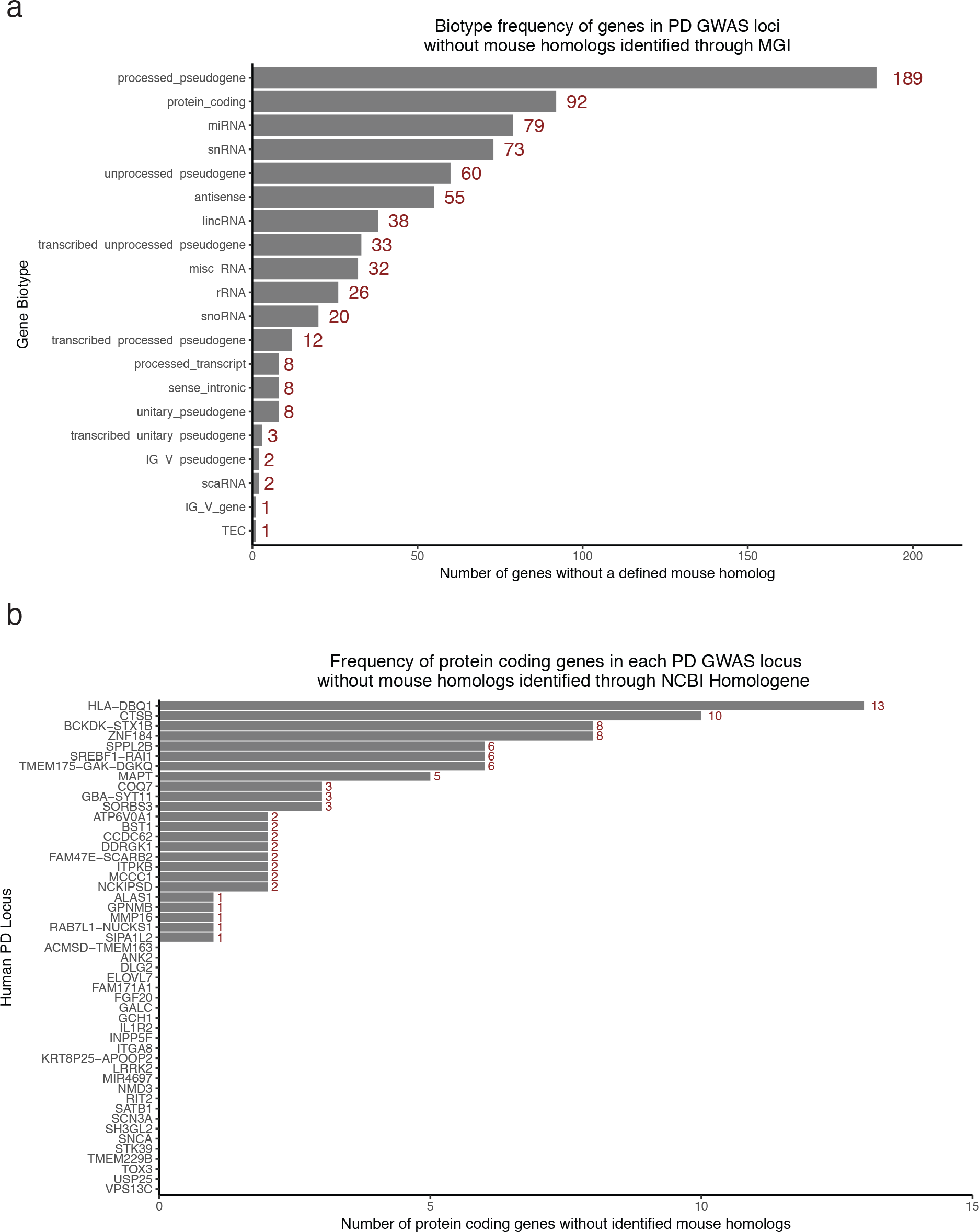
The distribution of gene biotypes assigned to genes extracted from PD GWAS loci. Figure 4 - supplement 1. The distribution of gene biotypes assigned to genes extracted from PD GWAS loci. a) Barplot displaying the frequency of gene biotypes in the 742 genes without mouse homologs identified in PD GWAS loci. Only 92/742 of those genes are annotated as protein coding. b) Barplot displaying the frequency of protein coding genes without mouse homologs in each PD GWAS locus studied. 24 loci include at least one protein coding gene without a mouse homolog.

**Table 1.**
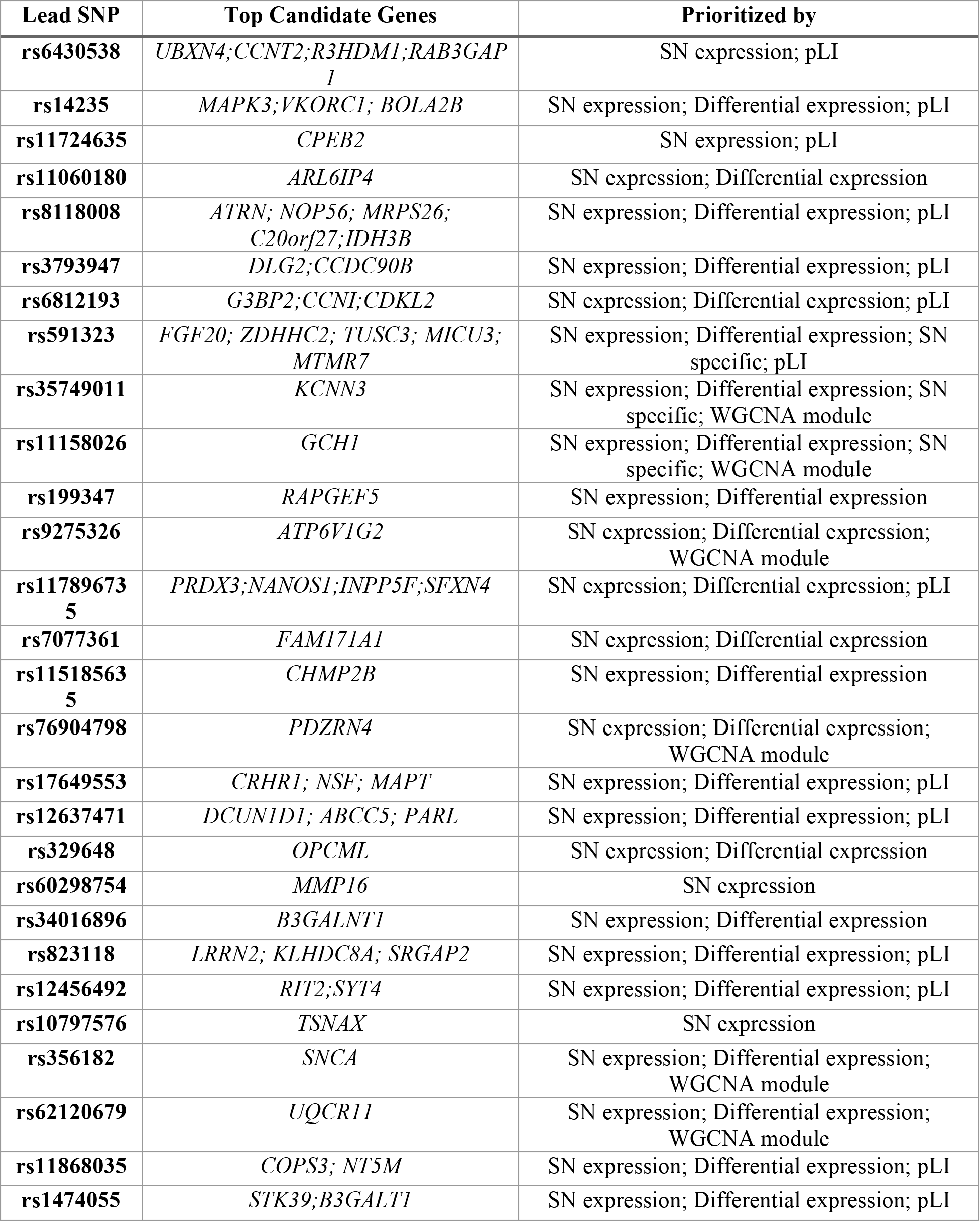
Summary of the systematic scoring of genes in 49 GWAS loci associated with PD. Scoring was carried out at described in the Results and Methods. Candidate genes are presented for each of 49 PD GWAS loci analyzed. Information for each PD GWAS locus is presented including the lead SNP for each locus, the sprioritized genes in each locus, and which data prioritized the top genes. Detailed scoring for each gene can be found in Supplementary File 9.

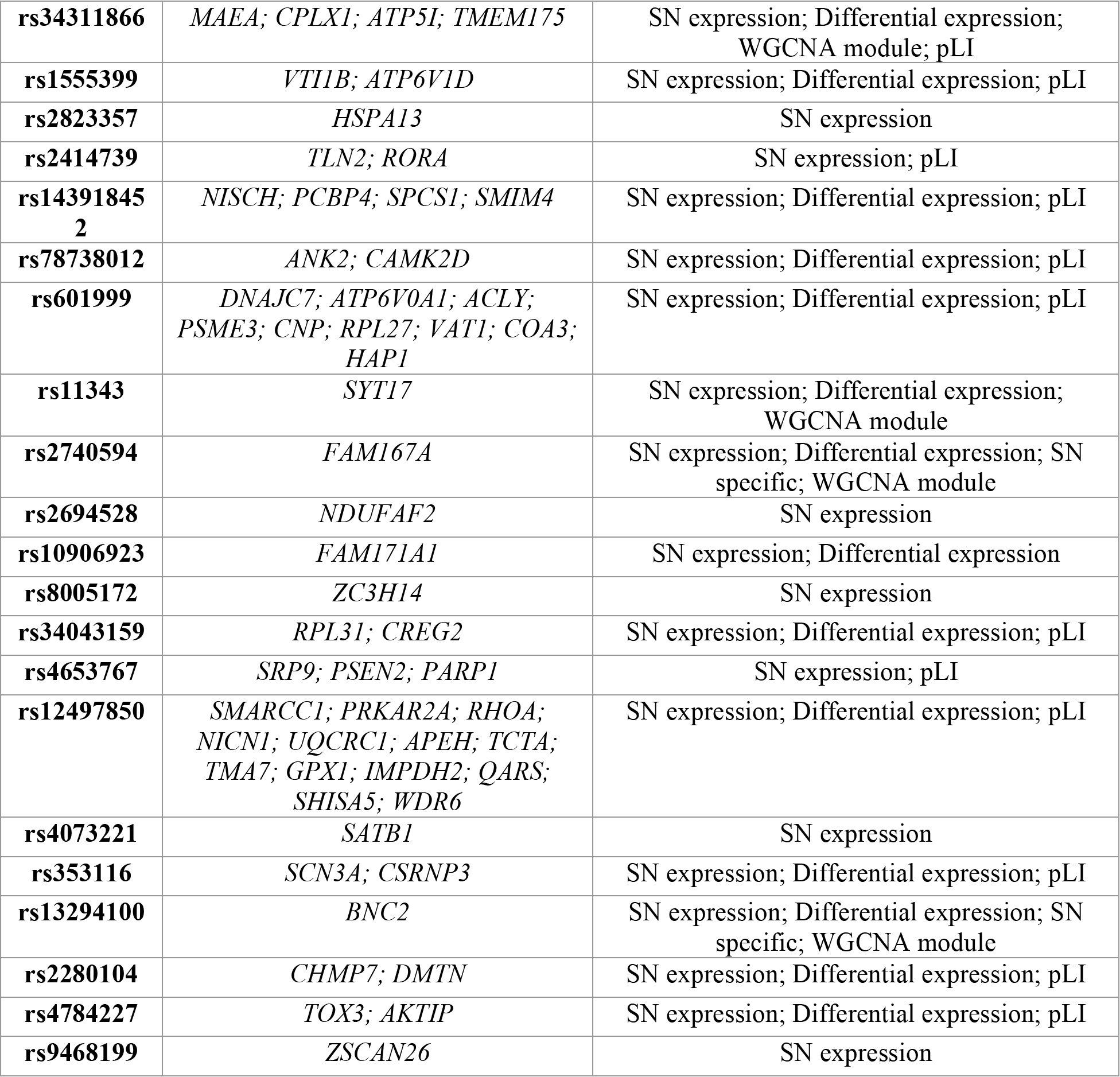

In order to prioritize likely diseases-associated genes in the remaining 46 loci, we scored genes on three criteria: whether genes were identified as specific markers for the P7.MB.4 (SN) cluster (Supplementary File 2), whether the genes were differentially expressed between all P7 DA neuron populations, and whether the genes were included in PD gene set enriched and SN correlated gene modules uncovered in WGCNA (Supplementary File 6). This strategy facilitated further prioritization of a single gene in 22 additional loci including *SNCA*, *LRRK2*, and *GCH1* loci (Figure 4a; Table 1; Supplementary File 9). Importantly, using this approach we indict the familial PD gene encoding alpha-synuclein (*SNCA*), as responsible for the observed PD association with rs356182 (Figure 4a, Table 1, Supplementary File 9). Thus, by using context-specific data alone, we were able to prioritize a single candidate gene in roughly half (~49%) of PD-GWAS associated loci.

Furthermore, at loci in which a single gene did not emerge, we identified dosage sensitive genes by considering the probability of being loss-of-function (LoF) intolerant (pLI) metric from the ExAC database^57,58^. Since most GWAS variation is predicted to impact regulatory DNA and in turn impact gene expression, it follows that genes in GWAS loci that are more sensitive to dosage levels may be more likely to be candidate genes. With that in mind, the pLI for each gene was used to further “rank” the genes within loci where a single gene was not prioritized. For those loci, including *MAPT* and *DDRGK1* loci (Figure 4a), we report a group of top scoring candidate genes (Table 1, Supplementary File 9). Expression of prioritized genes in the adult SN adds to the validity of the genes identified as possible candidates (Figure 4b).

Two interesting examples that emerge from this scoring are found at the *MAPT* and *TMEM175*-*GAK*-*DGKQ* loci. Although *MAPT* has previously been implicated in multiple neurodegenerative phenotypes, including PD (OMIM: 168600), we instead prioritize two genes before it (*CRHR1* and *NSF*; Table 1). We detect *Mapt* and *Nsf* expression consistently across all assayed DA neurons (Figure 4c). By contrast, expression of *Crhr1*, encoding the corticotropin releasing hormone receptor 1, is restricted to P7 DA neurons in the SN and the more mature OB neuronal populations (Figure 4c). Similarly, at the *TMEM175*-*GAK*-*DGKQ* locus, our data shows that although all three proximal genes are expressed in the SN, the adjacent *CPLX1* was one of the prioritized genes (Table 1, Supplementary File 9).

There are multiple lines of evidence that strengthen *CPLX1* as a candidate gene. Expression of CPLX1 is elevated both in the brains of PD patients and the brains of mice overexpressing the *SNCA* A53T PD mutation^59,60^. Additionally, mice deficient in *CPLX1* display an early-onset, cerebellar ataxia along with prolonged motor and behavioral phenotypes^61,62^. However, the impact of *Cplx1* deficiency on the integrity of the nigrostriatal pathway, to date, has not been explored. In order to confirm *CPLX1* as a candidate gene, we performed immunohistochemistry (IHC) for *Th* in the *Cplx1* knockout mouse model (Supplementary File 10, Supplementary File 11)^61–63^. We measured the density of *Th*+ innervation in the striatum of *Cplx1* -/- mice and controls (Figure 4d, Supplementary File 11) and found that *Cplx1* -/- mice had significantly lower Th+ staining in the striatum (p-value = 3.385e-08; Figure 4e). This indicates that *Cplx1* KO mice have less Th+ fiber innervation and a compromised nigrostriatal pathway, supporting its biological significance in MB DA populations and to PD.

The systematic identification of causal genes underlying GWAS signals is essential in order for the scientific and medical communities to take full advantage of all the GWAS data published over the last decade. Taken collectively, we demonstrate how scRNA-seq data from disease-relevant populations can be leveraged to illuminate GWAS results, facilitate systematic prioritization of GWAS loci implicated in PD, and can leads to the functional characterization of previously underexplored candidate genes.

## DISCUSSION

Midbrain DA neurons in the SN have been the subject of intense research since being definitively linked to PD nearly 100 years ago^64^. While degeneration of SN DA neurons in PD is well established, they represent only a subset of brain DA populations. It remains unknown why nigral DA neurons are particularly vulnerable. We set out to explore this question using scRNA-seq. Recently, others have used scRNA-seq to characterize the mouse MB, including DA neurons^34^. Here, we extend these data significantly, extensively characterizing the transcriptomes of multiple brain DA populations longitudinally and discovering GRNs associated with specific populations.

Most importantly, our data facilitate the iterative and biologically informed prioritization of gene candidates for all PD-associated genomic intervals. In practice, the gene closest to the lead SNP identified within a GWAS locus is frequently treated as the prime candidate gene, often without considering tissue-dependent context. Our study overcomes this by integrating genomic data derived from specific cell contexts with analyses that are agnostic to one another. We posit that genes pertinent to PD are likely expressed within SN DA neurons. This hypothesis is consistent with the recent description of the “omnigenic” nature of common disease, wherein variation impacting genes expressed in a disease tissue explain the vast majority of risk^7^.

First, we identify intervals that reveal one primary candidate, i.e. those that harbor only one SN-expressed gene. Next, we examine those intervals with many candidates, and prioritize based on a cumulative body of biological evidence. In total, we prioritize 5 or fewer candidates in 47/49 ( ~96%) PD GWAS loci studied, identifying a single gene in twenty-four loci (24/49; ~49%) and three or fewer genes in ~84% of loci (41/49). Ultimately this prioritization reduces the candidate gene list for PD GWAS loci dramatically from 1751 genes to 111 genes.

The top genes we identify in three PD loci (*SNCA*, *FGF20*, *GCH1*) have been directly associated with PD, MB DA development, and MB DA function^35^ (OMIM: 163890, 128230). Furthermore, our prioritization of *CPLX1* over other candidates in the *TMEM175*-*GAK*-*DGKQ* locus is supported by multiple lines of evidence. Additionally, we demonstrate that the integrity of the nigrostriatal pathway is disrupted in *Cplx1* knockout mice. Dysregulation of *CPLX1* RNA is also a biomarker in individuals with pre-PD prodromal phenotypes harboring the *PARK4* mutation (*SNCA* gene duplication)^65^. These results validate our approach and strengthen the argument for the use of context specific data in pinpointing candidate genes in GWAS loci.

Many of the genes prioritized (Table 1) have been shown to have various mitochondrial functions^66–72^. The identification of genes associated with mitochondrial functions is especially interesting in light of the “omnigenic” hypothesis of complex traits^7^. Since mitochondrial dysfunction has been extensively implicated in PD^73^, the prioritized genes may represent “core” genes that in turn can affect the larger mitochondrial-associated regulatory networks active in the disease relevant cell-type (SN DA neurons). It is notable that one of these genes is the presenilin associated rhomboid like gene or *PARL*. *PARL* cleaves *PINK1*, a gene extensively implicated in PD pathology and recently a variant in *PARL* has been associated with early-onset PD (OMIM: 607858)^74–76^.

While our method successfully prioritized one familial PD gene (*SNCA*), we do not prioritize *LRRK2*, another familial PD gene harbored within a PD GWAS locus. *Lrrk2* is not prioritized simply because it is not detectably expressed in our SN DA neuronal population. This is expected as numerous studies have reported little to no *Lrrk2* expression in *Th*+MB DA neurons both in mice and humans^77,78^. Instead, our method prioritizes *PDZRN4*. This result does not necessarily argue against the potential relevance of *LRRK2* but instead provides an additional candidate that may contribute to PD susceptibility. The same logic should be noted for two other PD-associated loci, wherein our scoring prioritizes different genes (*KCNN3* and *CRHR1*/*NSF*, respectively) than one previously implicated in PD (*GBA* and *MAPT*) (OMIM: 168600). Notably, *KCNN3*, *CRHR1*, and *NSF*, all have previous biological evidence making them plausible candidates^79–81^.

Studying disease-relevant tissue has proven to be essential for elucidating the genetic architecture underlying GWA signals^2^; our scoring method relies upon data from the most relevant cell-type to PD, SN DA neurons. While this study was under consideration for publication, Chang and colleagues^12^ endeavored to prioritize PD GWAS loci using publically available data. Although their pipeline strives to be “neuro-centric,” it is not predicated on the biological relevance of candidates to SN DA neurons.

Through comparison of the two scoring paradigms, the methods agree on at least one gene in 17/44 (~39%) jointly scored loci, including *SNCA* (Supplementary File 12), bolstering the evidence for those candidate genes. However, we see ~44% (31/71) of the genes prioritized by Chang, *et al*, are not expressed in either of the SN DA expression data sets used in our scoring scheme (Supplementary File 12), including *LRRK2* (addressed above). One prime example of this discrepancy is the *MCCC1* locus. Chang, *et al*, identify the *MCCC1* gene to be the prime candidate gene in the locus. However, we find that *MCCC1* is not expressed in SN DA neurons (Supplementary File 8). Instead, we prioritize *PARL*, a gene with an established role in PD pathogenesis^74–76^.

Our focus on disease relevant cell-type data also leads us to identify genes previously implicated in neurodegeneration, which make obvious candidates. For example, in the *TMEM175*-*GAK*-*DGKQ* locus, we identify *CPLX1* and functionally confirm its relevance. We also identify *ATRN* (attractin) as one of the candidate genes in the *DDRGK1* locus. Loss of *Atrn* has been shown to cause age-related neurodegeneration of SN DA neurons in rats^82,83^, making it an ideal candidate in the *DDRGK1* locus. Neither gene is identified using other metrics^12^ (Supplementary File 12).

Despite this success, we acknowledge several notable caveats. First, not all genes in PD-associated human loci have identified mouse homologs. Thus, it remains possible that we may have overlooked the contribution of some genes whose biology is not comprehensively queried in this study. Secondly, we assume that identified genetic variation acts in a manner that is at least preferential, if not exclusive, to SN DA neurons. Lastly, by prioritizing SN-expressed genes, we assume that PD variation affects genes whose expression in the SN does not require insult/stress. These caveats notwithstanding, our strategy sets the stage for a new generation of independent and combinatorial functional evaluation of gene candidates for PD-associated genomic intervals.

## MATERIALS AND METHODS

### Data availability

Raw data will be made available on Sequence Read Archive (SRA) and Gene Expression Omnibus (GEO) prior to publication. Summary data is available where code is available below (https://github.com/pwh124/DA_scRNA-seq).

### Code Availability

Code for analysis, for the production of figures, and summary data is deposited at https://github.com/pwh124/DA_scRNA-seq

### Animals

The Th:EGFP BAC transgenic mice (Tg(Th-EGFP)DJ76Gsat/Mmnc) used in this study were generated by the GENSAT Project and were purchased through the Mutant Mouse Resource & Research Centers (MMRRC) Repository (https://www.mmrrc.org/). Mice were maintained on a Swiss Webster (SW) background with female SW mice obtained from Charles River Laboratories (http://www.criver.com/). The Tg(Th-EGFP)DJ76Gsat/Mmnc line was primarily maintained through matings between Th:EGFP positive, hemizygous male mice and wild-type SW females (dams). Timed matings for cell isolation were similarly established between hemizygous male mice and wild-type SW females. The observation of a vaginal plug was defined as embryonic day 0.5 (E0.5). All work involving mice (husbandry, colony maintenance and euthanasia) were reviewed and pre-approved by the institutional care and use committee.

*Cplx1* knockout mice and wild type littermates used for immunocytochemistry were taken from a colony established in Cambridge using founder mice that were a kind gift of Drs K. Reim and N. Brose (Gottingen, Germany). *Cplx1* mice in this colony have been backcrossed onto a C57/Bl6J inbred background for at least 10 generations. All experimental procedures were licensed and undertaken in accordance with the regulations of the UK Animals (Scientific Procedures) Act 1986. Housing, rearing and genotyping of mice has been described in detail previously^61,62^. Mice were housed in hard-bottomed polypropylene experimental cages in groups of 5-10 mice in a housing facility was maintained at 21 −23°C with relative humidity of 55 ± 10%. Mice had *ad libitum* access to water and standard dry chow. Because homozygous knockout *Cplx1* mice have ataxia, they have difficulty in reaching the hard pellets in the food hopper and drinking from the water bottles. Lowered waterspouts were provided and access to normal laboratory chow was improved by providing mash (made by soaking 100 g of chow pellets in 230 ml water for 60 min until the pellets were soft and fully expanded) on the floor of the cage twice daily. *Cplx1* genotyping to identify mice with a homozygous or heterozygous deletion of the *Cplx1* gene was conducted as previously described^61^, using DNA prepared from tail biopsies.

### Dissection of E15.5 brains

At 15.5 days after the timed mating, pregnant dams were euthanized and the entire litter of embryonic day 15.5 (E15.5) embryos were dissected out of the mother and immediately placed in chilled Eagle’s Minimum Essential Media (EMEM). Individual embryos were then decapitated and heads were placed in fresh EMEM on ice. Embryonic brains were then removed and placed in Hank’s Balanced Salt Solution (HBSS) without Mg^2+^ and Ca^2+^ and manipulated while on ice. The brains were immediately observed under a fluorescent stereomicroscope and EGFP^+^ brains were selected. EGFP^+^ regions of interest in the forebrain (hypothalamus) and the midbrain were then dissected and placed in HBSS on ice. This process was repeated for each EGFP^+^ brain. Four EGFP^+^ brain regions for each region studied were pooled together for dissociation.

### Dissection of P7 brains

After matings, pregnant females were sorted into their own cages and checked daily for newly born pups. The morning the pups were born was considered day P0. Once the mice were aged to P7, all the mice from the litter were euthanized and the brains were then quickly dissected out of the mice and placed in HBSS without Mg^2+^ and Ca^2+^ on ice. As before, the brains were then observed under a fluorescent microscope, EGFP^+^ status for P7 mice was determined, and EGFP^+^ brains were retained. For each EGFP^+^ brain, the entire olfactory bulb was first resected and placed in HBSS on ice. Immediately thereafter, the EGFP^+^ forebrain and midbrain regions for each brain were resected and also placed in distinct containers of HBSS on ice. Five EGFP^+^ brain regions for each region were pooled together for dissociation.

### Generation of single cell suspensions from brain tissue

Resected brain tissues were dissociated using papain (Papain Dissociation System, Worthington Biochemical Corporation; Cat#: LK003150) following the trehalose-enhanced protocol reported by Saxena, et. al, 2012^84^ with the following modifications: The dissociation was carried out at 37°C in a sterile tissue culture cabinet. During dissociation, all tissues at all time points were triturated every 10 minutes using a sterile Pasteur pipette. For E15.5 tissues, this was continued for no more than 40 minutes. For P7, this was continued for up to 1.5 hours or until the tissue appeared to be completely dissociated.

Additionally, for P7 tissues, after dissociation but before cell sorting, the cell pellets were passed through a discontinuous density gradient in order to remove cell debris that could impede cell sorting. This gradient was adapted from the Worthington Papain Dissociation System kit. Briefly, after completion of dissociation according to the Saxena protocol^84^, the final cell pellet was resuspended in DNase dilute albumin-inhibitor solution, layered on top of 5 mL of albumin-inhibitor solution, and centrifuged at 70g for 6 minutes. The supernatant was then removed.

### FACS and single-cell collection

For each timepoint-region condition, pellets were resuspended in 200 μL of media without serum comprised of DMEM/F12 without phenol red, 5% trehalose (w/v), 25 μM AP-V, 100 μM kynurenic acid, and 10 μL of 40 U/μl RNase inhibitor (RNasin^®^ Plus RNase Inhibitor, Promega) at room temperature. The resuspended cells were then passed through a 40 uM filter and introduced into a Fluorescence Assisted Cell Sorting (FACS) machine (Beckman Coulter MoFlo Cell Sorter or Becton Dickinson FACSJazz). Viable cells were identified via propidium iodide staining, and individual neurons were sorted based on their fluorescence (EGFP+ intensity, See Figure 2 - supplement 2c) directly into lysis buffer in individual wells of 96-well plates for single-cell sequencing (2 μL Smart-Seq2 lysis buffer + RNAase inhibitor, 1 μL oligo-dT primer, and 1 μL dNTPs according to Picelli et al., 2014^85^. Blank wells were used as negative controls for each plate collected. Upon completion of a sort, the plates were briefly spun in a tabletop microcentrifuge and snap-frozen on dry ice. Single cell lysates were subsequently kept at −80°C until cDNA conversion.

### Single-cell RT, library prep, and sequencing

Library preparation and amplification of single-cell samples were performed using a modified version of the Smart-Seq2 protocol^85^. Briefly, 96-well plates of single cell lysates were thawed to 4°C, heated to 72°C for 3 minutes, then immediately placed on ice. Template switching first-strand cDNA synthesis was performed as described above using a 5’-biotinylated TSO oligo. cDNAs were amplified using 20 cycles of KAPA HiFi PCR and 5’-biotinylated ISPCR primer. Amplified cDNA was cleaned with a 1:1 ratio of Ampure XP beads and approximately 200 pg was used for a one-quarter standard sized Nextera XT tagmentation reaction. Tagmented fragments were amplified for 14 cycles and dual indexes were added to each well to uniquely label each library. Concentrations were assessed with Quant-iT PicoGreen dsDNA Reagent (Invitrogen) and samples were diluted to ~2 nM and pooled. Pooled libraries were sequenced on the Illumina HiSeq 2500 platform to a target mean depth of ~8.0 × 10^5^ 50bp paired-end fragments per cell at the Hopkins Genetics Research Core Facility.

### RNA sequencing and alignment

For all libraries, paired-end reads were aligned to the mouse reference genome (mm10) supplemented with the Th-EGFP^+^ transgene contig, using HISAT2^86^ with default parameters except: -p 8. Aligned reads from individual samples were quantified against a reference transcriptome (GENCODE vM8)^87^ supplemented with the addition of the eGFP transcript. Quantification was performed using cuffquant with default parameters and the following additional arguments: --no-update-check −p 8. Normalized expression estimates across all samples were obtained using cuffnorm^88^ with default parameters.

### Single-cell RNA data analysis

#### Expression estimates

Gene-level and isoform-level FPKM (Fragments Per Kilobase of transcript per Million) values produced by cuffquant^88^ and the normalized FPKM matrix from cuffnorm was used as input for the Monocle 2 single cell RNA-seq framework^89^ in R/Bioconductor^90^. Genes were annotated using the Gencode vM8 release^87^. A CellDataSet was then created using Monocle (v2.2.0)^89^ containing the gene FPKM table, gene annotations, and all available metadata for the sorted cells. All cells labeled as negative controls and empty wells were removed from the data. Relative FPKM values for each cell were converted to estimates of absolute mRNA counts per cell (RPC) using the Monocle 2 Census algorithm^17^ using the Monocle function “relative2abs.” After RPCs were inferred, a new cds was created using the estimated RNA copy numbers with the expression Family set to “negbinomial.size()” and a lower detection limit of 0.1 RPC.

#### QC Filtering

After expression estimates were inferred, the cds containing a total of 473 cells was run through Monocle’s “detectGenes” function with the minimum expression level set at 0.1 transcripts. The following filtering criteria were then imposed on the entire data set:

i. Number of expressed genes - The number of expressed genes detected in each cell in the dataset was plotted and the high and low expressed gene thresholds were set based on observations of each distribution. Only those cells that expressed between 2,000 and 10,000 genes were retained.
ii. Cell Mass - Cells were then filtered based on the total mass of RNA in the cells calculated by Monocle. Again, the total mass of the cell was plotted and mass thresholds were set based on observations from each distribution. Only those cells with a total cell mass between 100,000 and 1,300,000 fragments mapped were retained.
iii. Total RNA copies per cell - Cells were then filtered based on the total number of RNA transcripts estimated for each cell. Again, the total RNA copies per cell was plotted and RNA transcript thresholds were set based on observations from each distribution. Only those cells with a total mRNA count between 1,000 and 40,000 RPCs were retained.

A total of 410 individual cells passed these initial filters. Outliers found in subsequent, reiterative analyses described below were analyzed and removed resulting a final cell number of 396. The distributions for total mRNAs, total mass, and number of expressed, can be found in Figure 1 - supplement 1a-1c.

#### Log distribution QC

Analysis using Monocle relies on the assumption that the expression data being analyzed follows a log-normal distribution. Comparison to this distribution was performed after initial filtering prior to continuing with analysis and was observed to be well fit.

### Reiterative single-cell RNA data analysis

After initial filtering described above, the entire cds as well as subsets of the cds based on “age” and “region” of cells were created for recursive analysis. Regardless of how the data was subdivided, all data followed a similar downstream analysis workflow.

#### Determining number of cells expressing each gene

The genes to be analyzed for each iteration were filtered based on the number of cells that expressed each gene. Genes were retained if they were expressed in > 5% of the cells in the dataset being analyzed. These are termed “expressed_genes.” For example, when analyzing all cells collected together (n = 410), a gene had to be expressed in 20.5 cells (410 × 0.05 = 20.5) to be included in the analysis. Whereas when analyzing P7 MB cells (n = 80), a gene had to be expressed in just 4 cells (80 × 0.05 = 4). This was done to include genes that may define rare populations of cells that could be present in any given population.

#### Monocle model preparation

The data was prepared for Monocle analysis by retaining only the expressed genes that passed the filtering described above. Size factors were estimated using Monocle’s “estimateSizeFactors()” function. Dispersions were estimated using the “estimateDispersions()” function.

#### High variance gene selection

Genes that have a high biological coefficient of variation (BCV) were identified by first calculating the BCV by dividing the standard deviation of expression for each expressed gene by the mean expression of each expressed gene. A dispersion table was then extracted using the dispersionTable() function from Monocle. Genes with a mean expression > 0.5 transcripts and a “dispersion_empirical” >= 1.5*dispersion_fit or 2.0*dispersion_fit were identified as “high variance genes.”

#### Principal component analysis (PCA)

PCA was then run using the R “prcomp” function on the centered and scaled log2 expression values of the “high variance genes.” PC1 and PC2 were then visualized to scan the data for obvious outliers as well as bias in the PCs for age, region, or plates on which the cells were sequenced. If any visual outliers in the data was observed, those cells were removed from the original subsetted cds and all filtering steps above were repeated. Once there were no obvious visual outliers in PC1 or PC2, a screeplot was used plot the PCA results in order to determine the number of PCs that contributed most significantly to the variation in the data. This was manually determined by inspecting the screeplot and including only those PCs that occur before the leveling-off of the plot.

#### I. t-SNE and clustering

Once the number of significant PCs was determined, t-Distributed Stochastic Neighbor Embedding (t-SNE)^19^ was used to embed the significant PC dimensions in a 2-D space for visualization. This was done using the “tsne” package available through R with “whiten = FALSE.” The parameters “perplexity” and “max_iter” were tested with various values and set according what was deemed to give the cleanest clustering of the data.

After dimensionality reduction via t-SNE, the number of clusters was determined in an unbiased manner by fitting multiple Gaussian distributions over the 2D t-SNE projection coordinates using the R package ADPclust^91^ and the t-SNE plots were visualized using a custom R script. The number of genes expressed and the total mRNAs in each cluster were then compared.

### Differential expression Analyses

Since the greatest source of variation in the data was between ages (Figure 1), differential expression analyses and downstream analyses were performed separately for each age.

In order to find differentially expressed genes between brain DA populations at each age, the E15.5 and P7 datasets were annotated with regional cluster identity (“subset cluster”). Differential expression analysis was performed using the “differentialGeneTest” function from Monocle that uses a likelihood ratio test to compare a vector generalized additive model (VGAM) using a negative binomial family function to a reduced model in which one parameter of interest has been removed. In practice, the following models were fit: “~subset.cluster” for E15.5 or P7 dataset

Genes were called as significantly differentially expressed if they had a q-value (Benjamini-Hochberg corrected p-value) < 0.05.

### Cluster specific marker genes

In order to identify differentially expressed genes that were “specifically” expressed in a particular subset cluster, R code calculating the Jensen-Shannon based specificity score from the R package cummeRbund^92^ was used similar to what was described in Burns *et al*^93^.

Briefly, the mean RPC within each cluster for each expressed gene as well as the percentage of cells within each cluster that express each gene at a level > 1 transcript were calculated. The “.specificity” function from the cummRbund package was then used to calculate and identify the cluster with maximum specificity of each gene’s expression. Details of this specificity metric can be found in Molyneaux, *et al*^94^.

To identify subset cluster specific genes, the distribution of specificity scores for each subset cluster was plotted and a specificity cutoff was chosen so that only the “long right tail” of each distribution was included (i.e. genes with a specificity score above the cutoff chosen). For each iterative analysis, the same cutoff was used for each cluster or region (specificity ≥0.4). Once the specificity cutoff was chosen, genes were further filtered by only retaining genes that were expressed in >= 40% of cells within the subset cluster that the gene was determined to be specific for.

### Gene Set Enrichment Analyses

Gene set enrichment analyses were performed in two separate ways depending upon the situation. A Gene Set Enrichment Analysis (GSEA) PreRanked analysis was performed when a ranked list (e.g. genes ranked by PC1 loadings) using GSEA software available from the Broad Institute (v2.2.4)^95,96^. Ranked gene lists were uploaded to the GSEA software and a “GSEAPreRanked” analysis was performed with the following settings: ‘Number of Permutations’ = 1000, ‘Collapse dataset to gene symbols’ = true, ‘Chip platform(s)’ = GENE_SYMBOL.chip, and ‘Enrichment statistic’ = weighted. Analysis was performed against Gene Ontology (GO) collections from MSigDB, including c2.all.v5.2.symbols and c5.all.v5.2.symbols. Top ten gene sets were reported for each analysis (Supplementary File 1). Figures and tables displaying the results were produced using custom R scripts.

Unranked GSEA analyses for lists of genes was performed using hypergeometric tests from the R package clusterProfiler implemented through the functions ‘enrichGO’, ‘enrichKEGG’, and ‘enrichPathway’ with ‘pvalueCutoff set at 0.01, 0.1, 0.1, respectively with default settings^97^. These functions were implemented through the ‘compareCluster’ function when analyzing WGCNA data.

### Weighted Gene Co-Expression Network Analysis (WGCNA)

WGCNA was performed in R using the WGCNA package (v1.51)^45,46^ following established pipelines laid out by the packages authors (see https://labs.genetics.ucla.edu/horvath/CoexpressionNetwork/Rpackages/WGCNA/) for moredetail. Briefly, an expression matrix for all P7 neurons containing all genes expressed in >= 20 cells (n = 12628) was used with expression counts in log2(Transcripts + 1). The data were initially clustered in order to identify and remove outliers (n = 1) to leave 223 total cells (Figure 3 - supplement 1a). The soft threshold (power) for WGCNA was then determined by calculating the scale free topology model fit for a range of powers (1:10, 12, 14, 16, 18, 20) using the WGCNA function “pickSoftThreshold()” setting the networkType = “signed”. A power of 10 was then chosen based on the leveling-off of the resulting scale independence plot above 0.8 (Figure 3 - supplement 1b). Network adjacency was then calculated using the WGCNA function “adjacency()” with the following settings: power = 10 and type = “signed.” Adjacency calculations were used to then calculate topological overlap using the WGCNA function “TOMsimilarity()” with the following settings: TOMtype = “signed.” Distance was then calculated by subtracting the topological overlap from 1. Hierarchical clustering was then performed on the distance matrix and modules were identified using the “cuttreeDynamic” function from the dynamicTreeCut package^98^ with the following settings: deepSplit = T; pamRespectsDendro = FALSE, and minClusterSize = 20. This analysis initially identified 18 modules. Eigengenes for each module were then calculated using the “moduleEigengenes()” function and each module was assigned a color. Two modules (“grey” and “turquoise”) were removed at this point. Turquoise was removed because it contained 11567 genes or all the genes that could not be grouped with another module. Grey was removed because it only contained 4 genes, falling below the minimum set module size of 20. The remaining 16 modules were clustered (Figure 3 - supplement 1c) and the correlation between module eigengenes and subset cluster identity was calculated using custom R scripts. Significance of correlation was determined by calculated the Student asymptotic p-value for correlations by using the WGCNA “corPvalueStudent()” function. Gene set enrichments for modules were determined by using the clusterProfiler R package^97^. The correlation between the t-SNE position of a cell and the module eigengenes was calculated using custom R scripts.

### Prioritizing Genes in PD GWAS Loci

#### Topologically Associated Domain (TAD) and Megabase Gene Data

The data for human TAD boundaries were obtained from human embryonic stem cell (hESC) Hi-C data^49^ and converted from human genome hg18 to hg38 using the liftOver tool from UCSC Genome Browser (http://genome.ucsc.edu/). PD GWAS SNP locations in hg38 were intersected with the TAD information to identify TADs containing a PD GWAS SNP. The data for +/- 1 megabase regions surrounding PD GWAS SNPs was obtained by taking PD GWAS SNP locations in hg38 and adding or subtracting 1e+06 from each location. All hg38 Ensembl (version 87) genes that fell within the TADs or megabase regions were then identified by using the biomaRt R package^99,100^. All genes were then annotated with PD locus and SNP information. Mouse homologs for all genes were identified using human to mouse homology data from Mouse Genome Informatics (MGI) (http://www.informatics.jax.org/downloads/reports/HOM_MouseHumanSequence.rpt; Date accessed: 07/07/2017). Homologs of protein coding genes that did not have a mouse homolog in the data above were manually curated by searching the human gene name in the MGI database (http://www.informatics.jax.org/). Of the 742 genes with no mouse homologs, 92 (92/742, ~12%) were annotated as protein coding genes (Figure 4 - supplement 1a). 24 loci include at least one protein coding gene with no identified, one-to-one mouse homolog (Figure 4 - supplement 1b). All 1009 genes with mouse homologs are annotated as “protein_coding.” Gene homologs were manually annotated using the MGI database if a homolog was found to exist. The TAD and megabase tables were then combined to create a final PD GWAS locus-gene table.

#### PD GWAS Loci Gene Scoring

Genes within PD GWAS loci were initially scored using two gene lists: Genes with an average expression ≥0.5 transcripts in the SN cluster in our data (points = 1) and genes with an average expression ≥0.5 transcripts in the SN population in La Manno, *et al*^34^ (points = 1). Further prioritization was accomplished by using three gene lists: genes that were differentially expressed between subset clusters (points = 1); Genes found to be “specifically” expressed in the P7 MB SN cluster (points = 1); Genes found in the WGCNA modules that are enriched for PD (points = 1). Expression in the SN cluster was considered the most important feature and was weighted as such through the use of two complementary datasets with genes found to be expressed in both receiving priority. Furthermore, a piece of external data, pLI scores for each gene from the ExAC database^58^, was added to the scores in order to rank loci that were left with ≥2 genes in the loci after the initial scoring. pLI scores (fordist_cleaned_exac_r03_march16_z_pli_rec_null_data.txt) were obtained from http://exac.broadinstitute.org/ (Date dowloaded: March 30, 2017).

### In situ hybridization

*In situ* hybridization data was downloaded from publically available data from the Allen Institute through the Allen Brain Atlas (http://www.brain-map.org/). The image used in Figure 3a was obtained from the Reference Atlas at the Allen Brain Atlas (http://mouse.brain-map.org/static/atlas). URLs for all Allen Brain Atlas in situ data analyzed and downloaded for SN marker genes (Figure 3b) are available in Supplementary File 6. Data for SN expression *in situ* data for PD GWAS genes (Figure 4b) were obtained from the following experiments: 1056 (*Th*), 79908848 (Snca), 297 (Crhr1), 74047915 (*Atp6v1d*), 72129224 (*Mmp16*), and 414 (*Cntn1*). Data accessed on 03/02/17.

### Single molecule in situ hybridization (smFISH)

For in *situ* hybridization experiments, untimed pregnant Swiss Webster mice were ordered from Charles River Laboratories (Crl:CFW(SW); http://www.criver.com/). Mice were maintained as previously described. Pups were considered P0 on the day of birth. At P7, the pups were decapitated, the brain was quickly removed, and the brain was then washed in 1× PBS. The intact brain was then transferred to a vial containing freshly prepared 4% PFA in 1× PBS and incubated at 4°C for 24 hours. After 24 hours, brains were removed from PFA and washed three times in 1× PBS. The brains were then placed in a vial with 10% sucrose at 4°C until the brains sunk to the bottom of the vial (usually ~1 hour). After sinking, brains were immediately placed in a vial containing 30% sucrose at 4° until once again sinking to the bottom of the vial (usually overnight). After cryoprotection, the brains were quickly frozen in optimal cutting temperature (O.C.T.) compound (Tissue-Tek) on dry ice and stored at −80°C until use. Brains were sectioned at a thickness of 14 micrometers and mounted on Superfrost Plus microscope slides (Fisherbrand, Cat. # 12-550-15) with two sections per slide. Sections were then dried at room temperature for at least 30 minutes and then stored at −80°C until use.

RNAscope *in situ* hybridization (https://acdbio.com/) was used to detect single RNA transcripts. RNAscope probes were used to detect *Th* (C1; Cat No. 317621, Lot: 17073A), *Slc6a3* (C2; Cat No. 315441-C2, Lot: 17044A), *Lhx9* (C3; Cat No. 495431-C3, Lot: 17044A), and *Ldb2* (C3; Cat No. 466061-C3, Lot: 17044A). The RNAscope Fluorescent Multiplex Detection kit (Cat No. 320851) and the associated protocol provided by the manufacturer were used. Briefly, frozen tissues were removed from −80° and equilibrated at room temperature for 5 minutes. Slides were then washed at room temperature in 1× PBS for 3 minutes with agitation. Slides were then immediately washed in 100% ethanol by moving the slides up and down 5-10 times. The slides were then allowed to dry at room temperature and hydrophobic barriers were drawn using a hydrophobic pen (ImmEdge Hydrophobic Barrier PAP Pen, Vector Laboratories, Cat. # H-4000) around the tissue sections. The hydrophobic barrier was allowed to dry overnight. After drying, the tissue sections were treated with RNAscope Protease IV at room temperature for 30 minutes and then slides were washed in 1× PBS. Approximately 100 uL of multiplex probe mixtures (C1 - *Th*, C2 - *Slc6a3*, and C3 - one of *Lhx9* or *Ldb2*) containing either approximately 96 uL C1: 2 uL C2: 2 uL C3 (*Th*:*Slc6a3*:*Lhx9*) or 96 uL C1: 0.6 uL C2: 2 uL C3 (*Th:Slc6a3:Ldb2*) were applied to appropriate sections. Both mixtures provided adequate *in situ* signals. Sections were then incubated at 40° for 2 hours in the ACD HybEZ oven. Sections were then sequentially treated with the RNAscope Multiplex Fluorescent Detection Reagents kit solutions AMP 1-FL, AMP 2-FL, AMP 3-FL, and AMP 4 Alt B-FL, with washing in between each incubation, according to manufacturer’s recommendations. Sections were then treated with DAPI provided with the RNAscope Multiplex Fluorescent Detection Reagents kit. One drop of Prolong Gold Antifade Mountant (Invitrogen, Cat # P36930) was then applied to each section and a coverslip was then placed on the slide. The slides were then stored in the dark at 4°C overnight before imaging. Slides were further stored at 4°C throughout imaging. Manufacturer provided positive and negative controls were also performed alongside experimental probe mixtures according to manufacturer’s protocols. Four sections that encompassed relevant populations in the P7 ventral MB (SN, VTA, etc.) were chosen for each combination of RNAscope smFISH probes and subsequent analyses.

### smFISH Confocal Microscopy

RNAscope fluorescent *in situ* experiments were analyzed using the Nikon A1 confocal system equipped with a Nikon Eclipse Ti inverted microscope running Nikon NIS-Elements AR 4.10.01 64-bit software. Images were captured using a Nikon Plan Apo *λ* 60×/1.40 oil immersion lens with a common pinhole size of 19.2 μM, a pixel dwell of 28.8 μs, and a pixel resolution of 1024 × 1024. DAPI, FITC, Cy3, and Cy5 channels were used to acquire RNAscope fluorescence. Positive and negative control slides using probe sets provided by the manufacturer were used in order to calibrate laser power, offset, and detector sensitivity, for all channels in all experiments performed.

### smFISH image analysis and processing

Confocal images were saved as .nd2 files. Images were then processed in ImageJ as follows. First, the .nd2 files were imported into ImageJ and images were rotated in order to reflect a ventral MB orientation with the ventral side of the tissue at the bottom edge. Next the LUT ranges were adjusted for the FITC (range: 0-2500), Cy3 (range: 0-2500), and Cy5 (range: 0-1500) channels. All analyzed images were set to the same LUT ranges. Next, the channels were split and merged back together to produce a “composite” image seen in Figure 2. Scale bars were then added. Cells of interest were then demarcated, duplicated, and the channels were split. These cells of interest were then displayed as the insets seen in Figure 2.

### Immunohistochemistry and quantification of *Th* striatum staining in *Cplxl* mice

Mice (N=8 Cplx 1^1-/-^; N=3 WT littermates; ages between 4-7.5 weeks) were euthanized and their brains fresh-frozen on powdered dry ice. Brains were sectioned at 35 mm and sections and mounted onto Superfrost-plus glass slides (VWR International, Poole, UK). Sections were peroxidase inactivated, and one in every 10 sections was processed immunohistochemically for tyrosine hydroxylase. Sections were incubated in primary anti-tyrosine hydroxylase antibody (AB152, Millipore) used at 1/2000 dilution in 1% normal goat serum in phosphate-buffered saline and 0.2% Triton X-100 overnight at 4°C. Antigens were visualised using a horseradish peroxidase-conjugated anti-rabbit second antibody (Vector, PI-1000, 1/2000 dilution) and visualized using diaminobenzidine (DAB; Sigma). The slides were stored in the dark (in black slide boxes) at room temperature (21 C).

Images of stained striatum were taken using a Nikon AZ100 microscope equipped with a 2x lens (Nikon AZ Plan Fluor, NA 0.2, WD45), a Nikon DS-Fi2 camera, and NIS-Elements AR 4.5 software. Appropriate zoom and light exposure were determined before imaging and kept constant for all slides and sections. Density of Th+ DAB staining was measured using ImageJ software. Briefly, images were imported into ImageJ and the background was subtracted (default 50 pixels with “light background” selected). Next, images were converted to 8-bit and the image was inverted. Five measurements of density were taken for each side of a striatum in a section along with a density measurement from adjacent, unstained cortex. Striosomes were avoided during measuring when possible. Striatal measurements had background (defined as staining in the adjacent cortex in a section) subtracted. The mean section measurements (intensity/pixels squared) for each brain were calculated and represented independent measurements of the same brain. Variances were compared between the WT and KO populations. A two sample t-test was then used to compare WT vs. *Cplx1* -/- section densities with the following parameters in R using the “t.test” function: alternative = “two-sided”, var.equal = “T”.

## ACKNOWLEDGEMENTS

The authors wish to thank Stephen M. Brown for implementation and optimization of smFISH. Dr. Zhiguang Zheng and Mrs. Wendy Leavens for excellent technical support with the *Cplx1* knockout mice and immunohistochemistry and Drs Kerstin Reim and Niels Brose for the gift of the founder mice for the Cambridge Cplx1 knockout mice colony. This research was supported in part by US National Institutes of Health grants R01 NS62972 and MH106522 to ASM and a grant from CHDI *Inc*. to AJM.

## AUTHOR CONTRIBUTIONS

PWH, ASM, and LAG designed the study and wrote the paper. PWH, SAM, WDL, GAC, and AJM performed the experiments. PWH and LAG implemented the computational algorithms to process the raw data and conduct analyses thereof. PWH, LAG, and ASM analyzed and interpreted the resulting data. LAG contributed novel computational pipeline development. Correspondence to ASM and LAG.

## Supplemental File Descriptions

Supplementary File 1. A table with gene set enrichment analysis (GSEA) results for outliers removed during iterative analyses.

Supplementary File 2. A table with marker genes found for all 13 identified DA neuron populations.

Supplementary File 3. A table summarizing marker genes and observations that led to the biological classification of all 13 DA neuron populations

Supplementary File 4. A table showing marker genes of SN DA neurons with previous literature evidence of marking the SN.

Supplementary File 5. A table showing novel marker genes of SN DA neurons with summary of SN expression for each from Allen Brain Atlas (ABA) *in situ* data.

Supplementary File 6. A table showing all genes that comprise each identified WGCNA module.

Supplementary File 7. A table with Gene Ontology, Reactome, and KEGG enrichment results for all WGCNA modules.

Supplementary File 8. A table with meta-data for each locus in Table 1. This includes the “Lead SNP” associated with each locus, the “Closest Genes” to the lead SNP, and whether or not the closest genes are expressed (“Closest Gene Expressed”). This also has meta-data for genes in each locus including: the number of human genes (“num_genes”), the number of genes expressed in either of the SN DA scRNA-seq datasets used in scoring (“num_expressed_either”), the number of genes expressed in both SN DA scRNA-seq datasets using in scoring (“num_expressed_both”), the number of genes that had a one-to-one mouse homolog (“num_homolog”), and the number of genes that did not have a one-to-one mouse homolog (“num_no_homolog”).

Supplementary File 9. A table with detailed prioritization scoring for all genes within PD GWAS loci.

Supplementary File 10. A table summarizing information about *Cplx1* and WT mice used in this study including mouse name, age, genotype, the number of striatal sections measured, and the date immunohistochemistry was performed.

Supplementary File 11. A table showing all measurements taken for *Cplx1* and WT mice.

Supplementary File 12. A table summarizing the comparison of PD GWAS gene prioritization metrics found in this paper and in Chang, *et al* (2017).

